# A RabGAP-Rab GTPase pair regulates plant autophagy and immunity

**DOI:** 10.1101/2023.07.03.547386

**Authors:** Enoch Lok Him Yuen, Alexandre Y Leary, Marion Clavel, Yasin Tumtas, Azadeh Mohseni, Lorenzo Picchianti, Mostafa Jamshidiha, Pooja Pandey, Cian Duggan, Ernesto Cota, Yasin Dagdas, Tolga O Bozkurt

## Abstract

Plants rely on autophagy and membrane trafficking to tolerate stress, combat infections, and maintain cellular homeostasis. However, the molecular interplay between autophagy and membrane trafficking is poorly understood. Using an AI-assisted approach, we identified Rab3GAP-like (Rab3GAPL) as an important membrane trafficking node that controls plant autophagy negatively. Rab3GAPL suppresses autophagy by binding to ATG8, the core autophagy adaptor, and deactivating Rab8a, a small GTPase essential for autophagosome formation and defense-related secretion. Rab3GAPL from *Nicotiana benthamiana*, but not its mutated form deficient in ATG8 binding, reduced autophagic flux in *N. benthamiana* and Arabidopsis. Furthermore, *Rab3GAPL*-knockout mutants of the liverwort *Marchantia polymorpha* exhibited enhanced autophagic flux under both normal and heat stress conditions, suggesting that Rab3GAPL’s negative regulatory role in autophagy is conserved in land plants. Beyond autophagy regulation, Rab3GAPL modulates focal immunity against the oomycete pathogen *Phytophthora infestans* by preventing defense-related secretion. Altogether, our results suggest that Rab3GAPL acts as a molecular rheostat to coordinate autophagic flux and defense-related secretion by restraining Rab8a-mediated trafficking. This unprecedented interplay between a RabGAP-Rab pair and ATG8 sheds new light on the intricate membrane transport mechanisms underlying plant autophagy and immunity.

## Introduction

Plants face various environmental stresses such as temperature fluctuations, drought, and nutrient deficiencies on a daily basis. To effectively cope with such challenges and thrive in diverse environments, plants rely on autophagy, a catabolic process that aids in maintaining cellular homeostasis (*1, 2*). Autophagy facilitates the degradation of unwanted or harmful cellular components via the lytic compartments of the cells known as lysosomes or vacuoles. Importantly, autophagy also plays a vital role in plant immunity, although the specific underlying mechanisms are still under debate (*3*). For instance, autophagy can sequester pathogen molecules and even viruses for degradation (*4, 5*). On the contrary, plant pathogens have evolved strategies to evade or manipulate antimicrobial autophagy, underscoring its significance in plant defense (*6*).

Autophagy is a multistep process initiated by the induction of an isolation membrane that expands and closes, forming the mature autophagosomes — atypical vesicles with double membranes (*7*). The process of autophagy is orchestrated by a set of highly conserved autophagy-related (ATG) proteins that coordinate the biogenesis and maturation of autophagosomes (*8*). Studies in the last decade have revealed that autophagy is not only a bulk degradation process that is triggered during starvation, but also encompasses selective pathways that recycle specific cellular components via dedicated cargo receptors, adaptors, or modulators (*9*). At the heart of the autophagy machinery lies the ubiquitin-like protein ATG8, which functions as a hub to recruit autophagy cargo receptors and modulatory proteins. Once ATG8 undergoes lipid modification by the autophagy initiation complex, it becomes embedded within the inner and outer leaflets of the autophagosomal membranes. This localization of lipidated ATG8 is pivotal in coordinating the formation, transport, and fusion events of autophagosomes (*10, 11*). ATG8-interacting proteins contain short linear motifs termed ATG8-interacting motifs (AIMs, also known as LC3-interacting regions (LIRs) (*12*)). The canonical AIM sequence ([W,Y,F][X][X][L,I,V]) consists of an aromatic amino acid followed by any two amino acids and a hydrophobic residue, which are docked onto the W and L pockets on ATG8 (*13*). The discovery of proteins that carry functional AIMs is crucial for elucidating various aspects of autophagy. We have recently developed an AI-guided pipeline, utilizing Alphafold2-multimer (AF2-M), for prediction of both canonical and non-canonical AIM residues, enabling fast-forward discovery of autophagy receptors and modulators (*14*).

Intriguingly, pathogens can also exploit ATG8 as a hub to either subvert antimicrobial autophagy or tap into nutrient sources of their host cells (*15, 16*). Previously, we have shown that the Irish potato famine pathogen *Phytophthora infestans* secretes the effector protein PexRD54 which carries a canonical AIM that is required to subvert defense-related autophagy (*17, 18*). PexRD54 also promotes autophagosome formation by mimicking starvation conditions. These pathogen-induced autophagic vesicles are subsequently diverted to the host-pathogen interface, possibly as nutrient resources (*15*). Notably, PexRD54 acts as a scaffold between ATG8 compartments and vesicles labeled by the small GTPase Rab8a, likely to channel host lipid sources to stimulate autophagosome biogenesis (*15*). This further underscores the dynamic relationship between vesicle trafficking and autophagy.

Rab GTPases (Rabs), key components that regulate vesicle formation, transport, tethering and fusion events, have been identified to participate in different stages of autophagy (*19–21*). For instance, yeast and plant Rab1 members are crucial for early autophagosome formation (*22, 23*), while mammalian and plant Rab8a members have also been implicated in autophagy (*15, 24*). Rabs function as molecular switches that dynamically transit between an active GTP-bound state and an inactive GDP-bound state. These switches are tightly regulated by guanine nucleotide exchange factors (GEFs) that promote GTP binding and GTPase-activating proteins (GAPs) that catalyze GTP hydrolysis. RabGAPs, in particular, deactivate their Rab substrates, thereby determining their localization and function (*25*). Recently, mammalian TBC (Tre2/Bub2/Cdc16) domain-containing RabGAPs have been discovered to carry functional AIMs and modulate autophagy (*26*). However, the role of TBC-free RabGAPs in autophagy has not yet been demonstrated. Furthermore, the Rab substrate of RabGAPs, the trafficking pathways they govern, and the extent to which they regulate autophagy remain unknown in plants.

Here, we identified a TBC-free RabGAP protein, Rab3GAPL, as a key regulator of vesicle trafficking that interacts with ATG8 and Rab8a to suppress plant autophagy. Rab3GAPL also modulates the immune response against *P. infestans* by perturbing Rab8a vesicle dynamics and impairing defense-related secretion towards the pathogen interface. We uncovered an intricate interplay between a RabGAP protein, its Rab substrate, and the core autophagy receptor ATG8, underscoring their vital roles in regulating membrane trafficking processes essential for both autophagy and immunity.

## Results

### 1. A plant RabGAP, Rab3GAPL, directly interacts with ATG8CL through its C-terminal AIM

To uncover the roles of endomembrane trafficking in plant autophagy, we set out to identify vesicle transport regulators that associate with the autophagy machinery. We used our recently established Alphafold2-multimer (AF2-M)-assisted approach (*14*) to identify candidate trafficking components from our previous proteomics screen of ATG8CL interactors in the solanaceous model plant *Nicotiana benthamiana* (*27*). Through AF2-M-assisted re-analysis of the ATG8CL proteome, we identified a Rab GTPase-activating protein that carries specific domains to govern both vesicle trafficking and autophagy. Specifically, this RabGAP comprises the conserved Rab3GAP-like catalytic subunit at its core (Rab3GAPL hereafter), flanked by two helix-bundles and an N-terminal alpha-helix, alongside a C-terminal AIM (WTIV) that is predicted by AF2-M to bind to the AIM docking site on ATG8CL (Fig. 1A-B and S1A). The predicted structure reveals a stable interaction interface between Rab3GAPL AIM and ATG8CL AIM docking sites (Fig. 1B and S1A).

**Figure 1.**
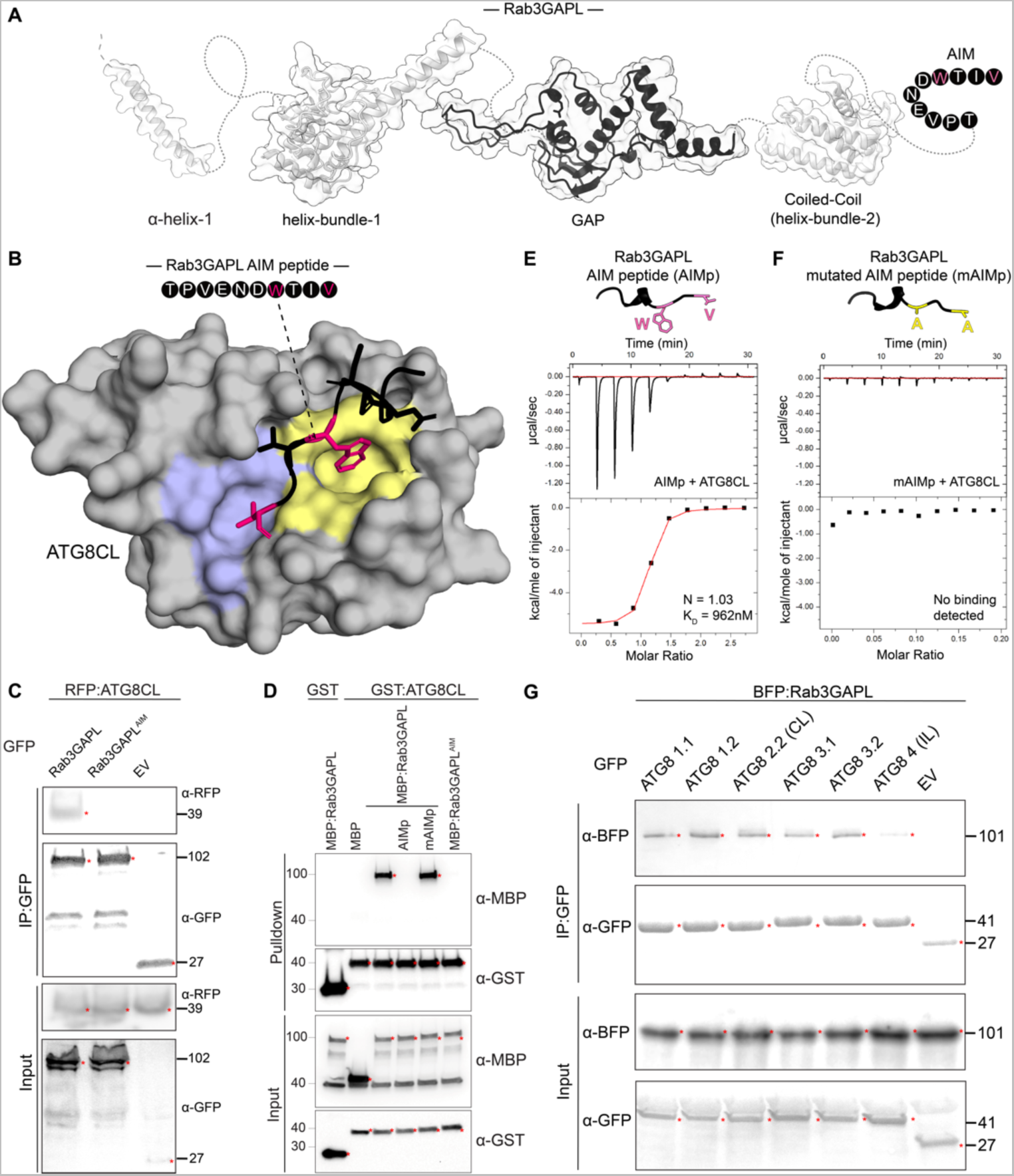
Rab3GAPL binds ATG8CL through a canonical AIM. (A) AF2 model of Rab3GAPL showing different regions (from N-terminal to C-terminal): A long alpha-helix (α-helix-1); a helix-bundle (helix- bundle-1) upstream of the conserved Rab3GAPL catalytic subunit; a coiled-coil region with a 4-helix bundle (helix-bundle-2), and a C-terminal AIM region. (B) AF2-M predicted structural model of Rab3GAPL and ATG8CL interaction displaying the docking of the Rab3GAPL-AIM peptide to the ATG8CL AIM pocket. Yellow and blue regions highlight W and L pockets on ATG8CL, respectively. (C) Rab3GAPL binds to ATG8CL via its AIM *in planta*. RFP:ATG8CL was transiently co-expressed with either GFP:Rab3GAPL, GFP:Rab3GAPL^AIM^, or GFP:EV. IPs were obtained with anti-GFP antiserum. Total protein extracts were immunoblotted. Red asterisks indicate expected band sizes. (D) *In vitro* GST pull-down assay shows Rab3GAPL-ATG8CL physical interaction is AIM-dependent. Bacterial lysates containing recombinant proteins were mixed and pulled down with glutathione magnetic agarose beads. Input and bound proteins were visualized by immunoblotting with anti-GST and anti-MBP antibodies. A peptide derivative (AIMp) of the pathogen effector PexRD54’s AIM region depleted Rab3GAPL from ATG8CL complexes, whereas the mutated AIM peptide (mAIMp) had no effect. (E) The AIM peptide of Rab3GAPL (Rab3GAPL-AIMp) directly binds to ATG8CL *in vitro*. The binding affinity of Rab3GAPL-AIMp to ATG8CL was determined using isothermal titration calorimetry (ITC). The upper panel shows heat differences upon injection of Rab3GAPL-AIMp into ATG8CL and the bottom panel shows integrated heats of injection and the best fit line to a single site binding model using MicroCal Origin. K_D_ = 962 nM, N = 1.03. (F) No binding was detected between the mutated AIM peptide of Rab3GAPL (Rab3GAPL-mAIMp) and ATG8CL using ITC. (G) Rab3GAPL binds to different ATG8 isoforms that are transiently expressed in *N. benthamiana*. BFP:Rab3GAPL was transiently co-expressed with either GFP:ATG8 1.1, GFP:ATG8 1.2, GFP:ATG8 2.2 (CL), GFP:ATG8 3.1, GFP:ATG8 3.2, GFP:ATG8 4 (IL) or GFP:EV. IPs were obtained with anti-GFP antiserum. Total protein extracts were immunoblotted. Red asterisks indicate expected band sizes.

To confirm the AIM as the mediator of the interaction between Rab3GAPL and ATG8CL, we substituted the key AIM residues tryptophan and valine with alanines (WTIV > ATIA) and performed co-immunoprecipitation (co-IP) assays. We generated N-terminal green fluorescent protein (GFP) fusions of wild-type (WT) Rab3GAPL and its AIM mutant (Rab3GAPL^AIM^) and investigated their interaction with ATG8CL. In contrast to GFP:Rab3GAPL, neither the GFP:Rab3GAP^AIM^ mutant nor the GFP control were able to pull down RFP:ATG8CL from *N. benthamina* protein extracts in the co-IP experiments (Fig. 1C). This observation demonstrates the specificity of the interaction between Rab3GAPL and ATG8CL, which is dependent on the presence of the C-terminal AIM as predicted by AF2-M (Fig. 1B-C). Importantly, the loss of ATG8CL-Rab3GAPL^AIM^ interaction cannot be attributed to altered localization or reduced stability of the AIM mutant, given the comparable protein levels and cytoplasmic localization patterns of both GFP:Rab3GAP and GFP:Rab3GAP^AIM^ constructs (Fig. 1C and S1B). These findings validate that Rab3GAPL interacts with ATG8CL through its C-terminal AIM, indicating a potential physical interaction between the two proteins.

To further investigate the potential AIM-mediated physical interaction between Rab3GAPL and ATG8CL, we performed *in vitro* glutathione S-transferase (GST) pull-down assays using MBP fusions of Rab3GAPL and Rab3GAPL^AIM^ in combination with GST:ATG8CL or GST (negative control) expressed in *Escherichia coli*. Consistent with AF2-M predictions and *in planta* co-IP assays, GST:ATG8CL specifically pulled down Rab3GAPL but not its AIM mutant Rab3GAPL^AIM^. Moreover, Rab3GAPL–ATG8CL interaction was abolished in the presence of the AIM peptide (AIMp) derived from the pathogen effector PexRD54 that binds ATG8CL, but not with the mutated AIM peptide (mAIMp), providing further support for the AIM-mediated physical binding of Rab3GAPL and ATG8CL (Fig. 1D).

To strengthen our findings, we conducted isothermal titration calorimetry (ITC) assays using a synthetic Rab3GAPL AIM peptide (Rab3GAPL-AIMp), which consists of the last 10 amino acid residues of Rab3GAPL that contains the AIM region. The Rab3GAPL-AIMp bound to ATG8CL with high affinity and in a one-to-one ratio (*K*_D_ = 962 nM and N = 1.03 based on ITC) (Fig. 1E). In contrast, we did not detect any association between the mutated AIM peptide of Rab3GAPL and ATG8CL (Fig. 1F). These results conclusively show that Rab3GAP’s C-terminal AIM is both necessary and sufficient to directly bind ATG8CL.

In plants, ATG8 has diversified into multiple isoforms (ATG8A–I), forming distinct ATG8 clades that potentially coordinate different selective autophagy pathways (*27, 28*). Therefore, we next set out to determine the specificity of the binding between Rab3GAPL and other solanaceous ATG8 isoforms. In co-IP assays using plant extracts, we observed that Rab3GAPL interacts with various potato ATG8 members exhibiting comparable affinities. Notably, the interaction between Rab3GAPL and the ATG8IL isoform appeared relatively weaker (Fig. 1G). This finding suggests that Rab3GAPL may have a broader functional role in autophagy by interacting with multiple ATG8 isoforms. All in all, these results show Rab3GAPL binds to the core autophagy protein ATG8 via a canonical C-terminal AIM.

### 2. Rab3GAPL negatively regulates autophagy in an AIM and GAP-dependent manner

Having established the physical interaction of Rab3GAPL and ATG8CL, we next investigated the role of Rab3GAPL in autophagy. We first investigated the impact of Rab3GAPL on autophagic puncta using confocal laser scanning microscopy (CLSM). To this end, we imaged cells transiently expressing GFP:Rab3GAPL, the AIM mutant GFP:Rab3GAPL^AIM^, or a GFP control alongside the autophagosome marker RFP:ATG8CL and quantified autophagosome numbers. In cells expressing GFP:Rab3GAPL we observed a greater than two-fold reduction in RFP:ATG8CL puncta compared to cells expressing a GFP control. In contrast, GFP:Rab3GAP^AIM^-expressing cells did not show any significant difference in the relative amount of RFP:ATG8CL puncta (Fig. 2A-B). The observed decrease in the quantity of autophagic puncta caused by overexpression of Rab3GAPL, but not its AIM mutant, signals at the possibility that Rab3GAPL negatively regulates autophagy, which relies on its ability to bind ATG8.

**Figure 2.**
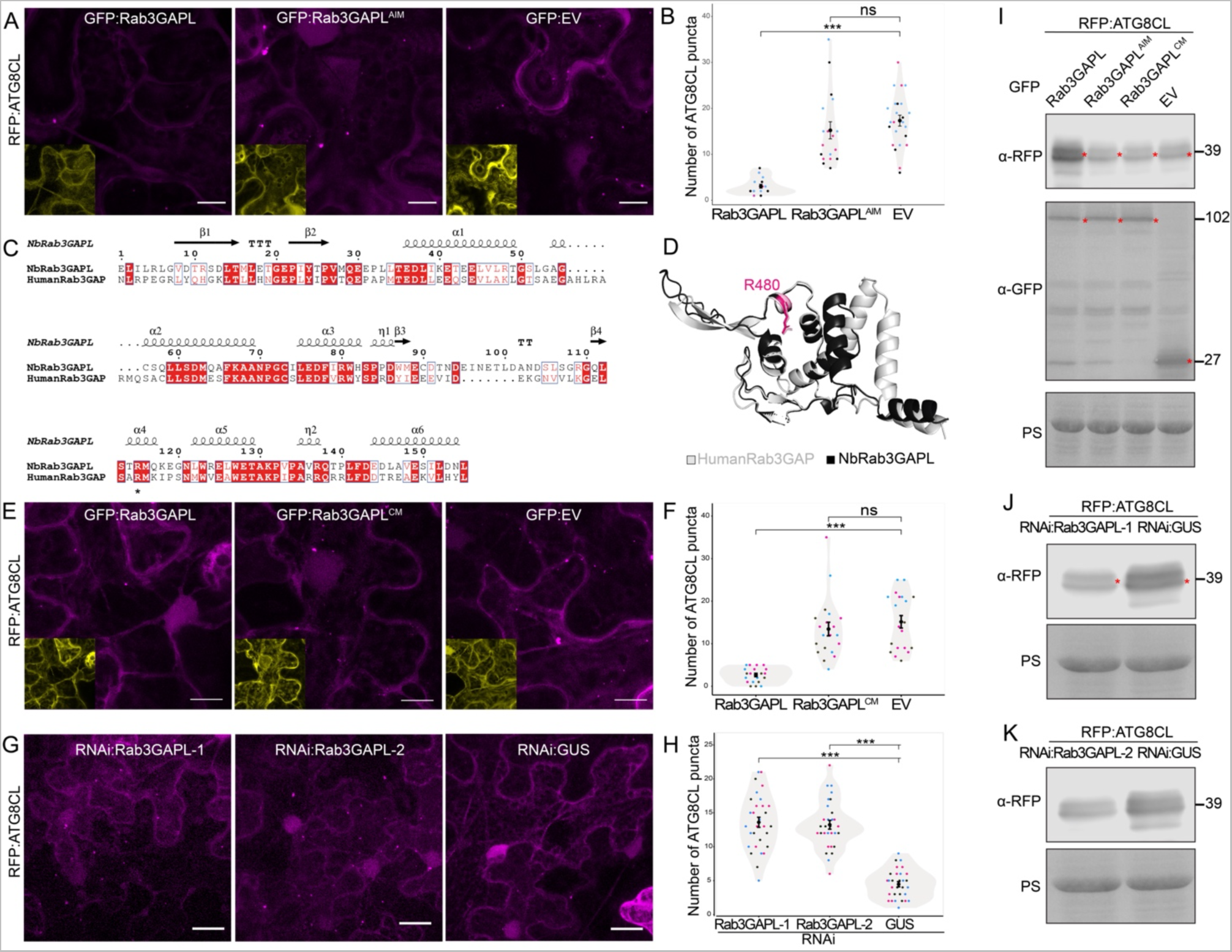
Rab3GAPL suppresses autophagy in an AIM and catalytic activity-dependent manner. (A-B) Rab3GAPL reduces the number of ATG8CL autophagosomes in an AIM-dependent manner. (A) Confocal micrographs of *N. benthamiana* leaf epidermal cells transiently expressing RFP:ATG8CL with GFP:Rab3GAPL, GFP:Rab3GAPL^AIM^ or GFP:EV. Images shown are maximal projections of 17 frames with 1.3 μm steps. Scale bars represent 10 μm. (B) Rab3GAPL expression significantly reduces ATG8CL autophagosomes (2, N = 18 images) compared to EV control (17, N = 18 images), while Rab3GAPL^AIM^ expression has no significant effect on the number of ATG8CL autophagosomes (13, N = 18 images) compared to EV control. Statistical differences were analyzed by Mann-Whitney U test in R. Measurements were highly significant when p<0.001 (***). (C) Amino acid alignment of the GAP domains of human Rab3GAP and *N. benthamiana* Rab3GAPL proteins. * denotes the conserved catalytic arginine finger. (D) Structural alignment of the GAP domains of human Rab3GAP and *N. benthamiana* Rab3GAPL. Structural predictions were obtained via AF2. The model shows conservation of the positioning of the catalytic arginine finger. (E-F) Rab3GAPL reduces the number of ATG8CL autophagosomes in a catalytic activity-dependent manner. (E) Confocal micrographs of *N. benthamiana* leaf epidermal cells transiently expressing RFP:ATG8CL with GFP:Rab3GAPL, GFP:Rab3GAPL^CM^ or GFP:EV. Images shown are maximal projections of 17 frames with 1.5 μm steps. Scale bars represent 10 μm. (F) Rab3GAPL expression significantly reduces ATG8CL autophagosomes (3, N = 20 images) compared to EV control (14.5, N = 20 images), while Rab3GAPL^CM^ expression has no significant effect on the number of ATG8CL autophagosomes (12, N = 20 images) compared to EV control. Statistical differences were analyzed by Mann-Whitney U test in R. Measurements were highly significant when p<0.001 (***). (G-H) RNAi-mediated silencing of Rab3GAPL increases the number of ATG8CL autophagosomes. (G) Confocal micrographs of *N. benthamiana* leaf epidermal cells transiently expressing RNAi:Rab3GAPL-1, RNAi:Rab3GAPL-2 or RNAi:GUS. Images shown are maximal projections of 24 frames with 1.3 μm steps. Scale bars represent 10 μm. (H) Silencing Rab3GAPL-1 (13, N = 30 images) or Rab3GAPL-2 (13, N = 30 images) significantly increases the number of ATG8CL autophagosomes compared to GUS silencing control (4.5, N = 30 images). Statistical differences were analyzed by Welch’s T-test in R. Measurements were highly significant when p<0.001 (***). (I-K) Rab3GAPL suppresses autophagic flux in an AIM and catalytic activity-dependent manner. (I) Western blot shows depletion of RFP:ATG8CL is reduced by GFP:Rab3GAPL compared to GFP:Rab3GAPL^AIM^, GFP:Rab3GAPL^CM^, or EV control. Total protein extracts were prepared 4 days post agroinfiltration and immunoblotted. Red asterisks show expected band sizes. (J-K) Western blots show depletion of RFP:ATG8CL is increased by silencing Rab3GAPL using either of the two silencing constructs (J) RNAi:Rab3GAPL-1, or (K) RNAi:Rab3GAPL-2 compared to the GUS silencing control. Total protein extracts were prepared 4 days post agroinfiltration and immunoblotted. Red asterisks show expected band sizes.

To determine the extent to which Rab3GAPL modulates autophagy, we next sought to determine whether the reduction in autophagosome numbers caused by Rab3GAPL overexpression is dependent on the GAP activity of Rab3GAPL. Previously, it was shown that the GAP function of human Rab3GAP was compromised by the mutation of the conserved arginine finger, which typically establishes connections with the γ-phosphate of the GTP nucleotide (*29*). The structural alignment of the Rab3GAPL and human Rab3GAP protein sequences revealed that the catalytic arginine finger in the human Rab3GAP is conserved in *N. benthamiana* (R480) with the consensus sequence of LSxRM (Fig. 2C). The AF2 predictions of the GAP domains of human and *N. benthamiana* Rab3GAPL showed a high level of structural conservation of the GAP domain architecture with a root-mean-square deviation (RMSD) value of 0.486. Additionally, the catalytic arginine finger was positioned consistently in both predicted structures (Fig. 2D and S2A). Based on these observations, we generated the catalytic mutant of the *N. benthamiana* Rab3GAPL (Rab3GAP^CM^ hereafter) by mutating the conserved arginine at position 480 to alanine (R480A). Unlike GFP:Rab3GAPL which reduces autophagosome numbers (Fig. 2A-B, E-F), GFP:Rab3GAPL^CM^ did not significantly alter the amount of RFP:ATG8CL-labeled autophagosomes compared to the GFP control (Fig. 2E-F). Comparing the AF2 models of Rab3GAPL and Rab3GAPCM did not reveal any global structural alterations resulting from the point mutation (Fig. S2B). To further ensure that Rab3GAPL^CM^ is stably expressed and that its overall structure is not disrupted, we tested its stability as well as its ability to associate with ATG8CL *in vivo*. In these assays, we also used a dual mutant of Rab3GAPL (Rab3GAPL^CM/AIM^) carrying both the AIM and GAP mutations as an additional control. Western blots of the plant protein extracts (input) and pull-down assays (output) showed that GFP:Rab3GAPL^CM^ was stably expressed and was able to interact with RFP:ATG8CL, whereas the negative control GFP:Rab3GAP^CM/AIM^ dual mutant did not associate with ATG8CL (Fig. S2C). These results substantiate structural predictions that the overall protein architecture of Rab3GAPL^CM^ is not perturbed. We conclude that the GAP activity of Rab3GAP is required for its ability to suppress ATG8CL- autophagosome numbers.

Next, we determined the effect of *Rab3GAPL* silencing on autophagy by quantifying the number of RFP:ATG8CL-autophagosomes. In contrast to the overexpression results, silencing *Rab3GAPL* with two independent hairpin RNA interference (RNAi) constructs—one targeting the coding region (RNAi:Rab3GAPL-1) and the second one targeting the three prime untranslated region (3’UTR) of *Rab3GAPL* (RNAi:Rab3GAPL-2)—increased the amount of RFP:ATG8CL puncta by greater than two-fold compared to a β-glucuronidase silencing control (RNAi:GUS) (Fig. 2G-H, S2D). These results suggest that Rab3GAPL suppresses autophagy via the AIM and GAP domains.

The potential reason for the reduction in autophagy puncta resulting from the overexpression of Rab3GAPL may be attributed to a decrease in autophagosome formation or an increase in autophagosome degradation. To address this, we conducted autophagic flux assays upon overexpression or silencing of Rab3GAPL. We first investigated the impact of Rab3GAPL overexpression on the protein levels of ATG8CL in *N. benthamiana*. To measure autophagic flux, we extracted proteins from *N. benthamiana* leaves co-expressing RFP:ATG8CL in combination with GFP:Rab3GAPL, GFP:Rab3GAPL^AIM^, GFP:Rab3GAPL^CM^ or an empty vector control at four days post transient expression and performed western blotting. In three independent experiments, overexpression of GFP:Rab3GAPL led to increased relative protein levels of RFP:ATG8CL, whereas overexpression of GFP:Rab3GAPL^AIM^, GFP:Rab3GAPL^CM^ or GFP vector control did not show the same effect (Fig. 2I and S2E). These results are in line with our findings that only Rab3GAPL, and not its AIM or GAP mutants, modifies the quantity of autophagosomes (Fig. 2A-B, E-F). In accordance with the overexpression assays, the attenuation of Rab3GAPL gene expression through RNAi:Rab3GAPL-1 and RNAi:Rab3GAPL-2 resulted in a decrease in RFP:ATG8CL levels relative to an RNA interference construct that targeted GUS (Fig. 2J-K and S2F). These results suggest that Rab3GAPL negatively regulates autophagy at the autophagosome biogenesis stage.

### 3. The negative regulatory role of Rab3GAPL in autophagy is broadly conserved in land plants

We next investigated whether the regulatory function of Rab3GAPL in autophagy is conserved in other plant species. To test this, we analyzed Rab3GAPL sequences from phylogenetically diverse plants, including wheat, Arabidopsis, *N. benthamiana*, potato, and the liverwort *Marchantia polymorpha*. Our analysis of Rab3GAPL sequences revealed that the GAP domain was conserved across all plant species (Fig. S3A). Likewise, the AIM is highly conserved among all tested plant species, with Arabidopsis being the exception. Arabidopsis Rab3GAPL has a deletion in the key AIM residue W, along with upstream negatively charged residues that are known to be essential for interactions with positively charged surface residues of the AIM pocket (Fig. 3A and S3A).

**Figure 3.**
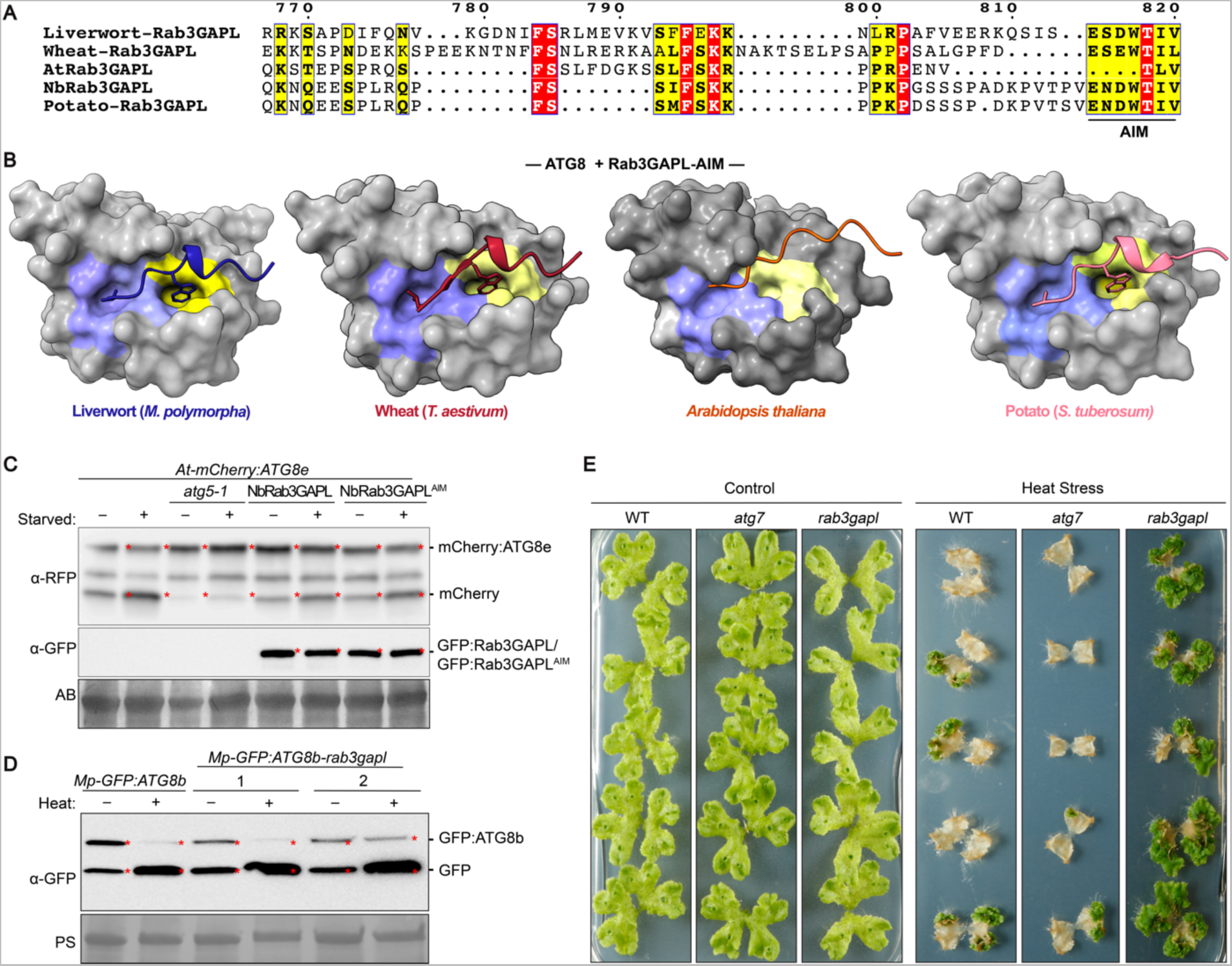
Rab3GAPL suppression of autophagic flux is widely conserved in land plants. (A). Pairwise sequence alignment comparisons of the N-terminals of Rab3GAPL in wheat, Arabidopsis, *N. benthamiana*, potato, and the liverwort *Marchantia polymorpha*. Alignments were obtained using the MUSCLE algorithm and were visualized and color-coded via ESPript 3.0 (*39*). The AIM is illustrated using a solid line. (B) AF2-M predictions of ATG8s with Rab3GAPL AIM sequences from liverwort (*M. polymorpha*), wheat (*Triticum aestivum*), *A. thaliana* and potato (*S. tuberosum*). Predicted models suggest AIM docking sites (W and L pockets, colored yellow and blue, respectively) on ATG8CL are associated with all tested AIMs except for the Arabidopsis Rab3GAPL AIM sequence. (C) *Arabidopsis thaliana* lines that overexpress Rab3GAPL have reduced ATG8 autophagic flux. Autophagic flux is measured as the ratio of free mCherry to full size mCherry:ATG8e. GFP:Rab3GAPL expression leads to reduced mCherry/mCherry:ATG8e protein signal ratio in both carbon starvation and control conditions compared to the control plants. Protein extracts were prepared using 6-day-old seedlings and immunoblotted. (D) *Marchantia polymorpha* Rab3GAPL-KO mutants have increased ATG8 autophagic flux. Autophagic flux analysis of WT and Rab3GAPL-KO mutants in MpEF::GFP:ATG8b background after 6 hours of heat stress treatment following 2 hours recovery. Flux is estimated as the ratio of free GFP to full size GFP:ATG8b. Both Rab3GAPL-KO mutants showed increased GFP/GFP:ATG8b protein signal ratio under heat stress and control conditions compared to the control plants. Protein extracts were prepared using 14-day-old thalli and immunoblotted. (E) *Mp-Rab3GAPL*-KO mutant showed enhanced recovery from heat stress compared to WT control plants. ATG7-KO plants were used as an autophagy- deficient control, which showed reduced recovery from heat stress compared to WT control plants. Transgenic plants were incubated either in normal condition (22°C) or heat stress condition (37°C).

The absence of the key AIM residues in Rab3GAPL of Arabidopsis (AtRab3GAPL) raised the question of whether it can bind to ATG8. To investigate this further, we used AF2-M to predict ATG8s in complex with Rab3GAPL sequences used in the MUSCLE analysis. Consistent with the multiple sequence alignment findings, the predicted models revealed that all tested Rab3GAPL proteins possess functional AIMs that occupy the AIM pockets on their respective ATG8 proteins, with the exception of AtRab3GAPL (Fig. 3B and S3B). These results suggest that while the regulatory function of Rab3GAPL in autophagy is largely conserved, it may not be present in certain plant species, including Arabidopsis.

To gain additional insights, we performed a BLAST search of the Rab3GAPL protein sequence against the Brassicales order of flowering plants, which includes Arabidopsis as well as economically important crops such as cabbage, broccoli, mustard, and papaya. Interestingly, while the papaya (Caricaceae) Rab3GAPL carries an intact AIM, Rab3GAPLs from the Brassicaceae and Cleomaceae families, which diverged over 40 million years ago (*30*), had deletions in their AIM residues as in the case of Arabidopsis (Fig. 3A and S3C). Considering our results that Rab3GAPL’s AIM residues are critical for autophagy suppression (Fig. 2), these findings suggest that the regulation of autophagy by Rab3GAPL may vary among different plant species. Further investigations are required to explore the impact of the loss of AIM in Rab3GAPLs from Brassicaceae and Cleomaceae.

Next, we investigated the potential of heterologous expression of Rab3GAPL from *N. benthamiana* (NbRab3GAPL) to inhibit autophagy in Arabidopsis. To assess this, we stably expressed GFP:NbRab3GAPL or GFP:NbRab3GAPL^AIM^ in Arabidopsis lines that express mCherry:ATG8e and measured the autophagic flux by analyzing the cleavage of mCherry (free mCherry) from mCherry:ATG8e fusion protein through western blotting (*31, 32*). We analyzed the impact of Rab3GAPL overexpression on basal autophagy and autophagy induced by carbon starvation by comparing the mCherry signal ratios in the GFP:Rab3GAPL/mCherry:ATG8e and GFP:Rab3GAPL^AIM^/mCherry:ATG8e lines, alongside the *atg5-1* autophagy deficient mutants that we used as a negative control. As expected, protein extracts from *atg5-1* lines showed lower mCherry/mCherry:ATG8e ratios in both carbon starvation and control conditions, indicating reduced autophagic flux across three independent experiments (Fig. 3C and S3D). Similarly, plants overexpressing GFP:Rab3GAPL exhibited reduced mCherry/mCherry:ATG8e ratios in both conditions, suggesting decreased autophagic degradation (Fig. 3C and S3D). In contrast, the GFP:Rab3GAPL^AIM^ mutant did not show any reduction in the autophagic degradation of mCherry:ATG8e, consistent with the autophagic flux assays in *N. benthamiana* (Fig. 2I-K). These results provide evidence that Rab3GAPL from *N. benthamiana* can suppress autophagy in Arabidopsis.

The conservation of the AIM and GAP domain in liverworts implies their significance in Rab3GAPL function throughout land plants, dating back at least 400 million years. Having observed the interference of Rab3GAPL with autophagic flux in two dicot models, we investigated whether the negative regulation of autophagy by Rab3GAPL is maintained in liverworts. To perform autophagic flux assays, we first generated two independent *M. polymorpha Rab3GAPL* CRISPR knockout mutants, designated as *Mp-rab3gapl-1* and *Mp-rab3gapl-2*, in a GFP:ATG8b background (Fig. S3E). We then compared the GFP/GFP:ATG8b protein signal ratios in the WT and mutant genotypes under both control and heat stress conditions across three independent experiments. The GFP/GFP:ATG8b ratios were consistently higher in the *Mp-rab3gapl* mutant lines than in the control plants, indicating increased ATG8 autophagic flux in *M. polymorpha* (Fig. 3D and S3F). These findings demonstrate that the knockout of *Rab3GAPL* enhances ATG8 autophagic flux in *M. polymorpha*.

Given the well-established beneficial impact of autophagy on heat stress tolerance and recovery (*33–36*), we next investigated whether enhanced autophagic flux observed in *Mp-rab3gapl* mutant lines can boost recovery from heat stress. To assess this, we compared the survival rates of WT plants, *Mp- rab3gapl* mutant lines, and *atg7* mutants deficient in autophagy functionality under heat stress conditions at 37°C, prior to recovery. We observed a notable recuperation from heat stress in the *Mp- rab3gapl* mutant plants compared to WT plants (Fig. 3E). In contrast, the autophagy-deficient *atg7* control lines exhibited reduced recovery from heat stress (Fig. 3E). These results are consistent with the studies which showed genetic interference of selective autophagy receptors lead to compromised heat tolerance due to the accumulation of protein aggregates that were highly ubiquitinated under heat stress (*34, 35, 37, 38*). However, further evidence is needed to conclude that enhanced recovery from heat stress in *M. polymorpha* is caused by enhanced autophagic activity in the absence of Rab3GAPL. Nevertheless, our results align with the beneficial impact of autophagy in stress tolerance and the role of Rab3GAPL as a negative regulator of autophagy conserved across different plant lineages.

### 4. Rab8a, a GTPase implicated in autophagy and immunity, is a substrate of Rab3GAPL

We next sought to identify the Rab GTPase partner of Rab3GAPL in autophagy regulation. We tested the interaction of Rab3GAPL with a panel of candidate Rabs from solanaceous plants—Rab1, Rab2, Rab8a, Rab8b—identified from our earlier autophagy interactome studies (*15, 17, 27*). As an additional control, we also included a *N. benthamiana* Rab11 member in the interaction assays, as mammalian Rab11 has been implicated in autophagy (*40*). The results from our co-IP assays indicate that Rab3GAPL strongly interacts with Rab8 members and weakly with Rab2. However, we did not observe any association between Rab3GAPL and Rab1, Rab11, or the GFP vector control (Fig. 4A). These results suggest that Rab8 members are candidate substrates of Rab3GAPL in autophagy regulation. Consistent with this notion, we have previously shown that Rab8a associates with ATG8CL and positively regulates autophagy by potentially facilitating the transport of lipids to the phagophore assembly sites (PAS) required for autophagosome biogenesis (*15*).

**Figure 4.**
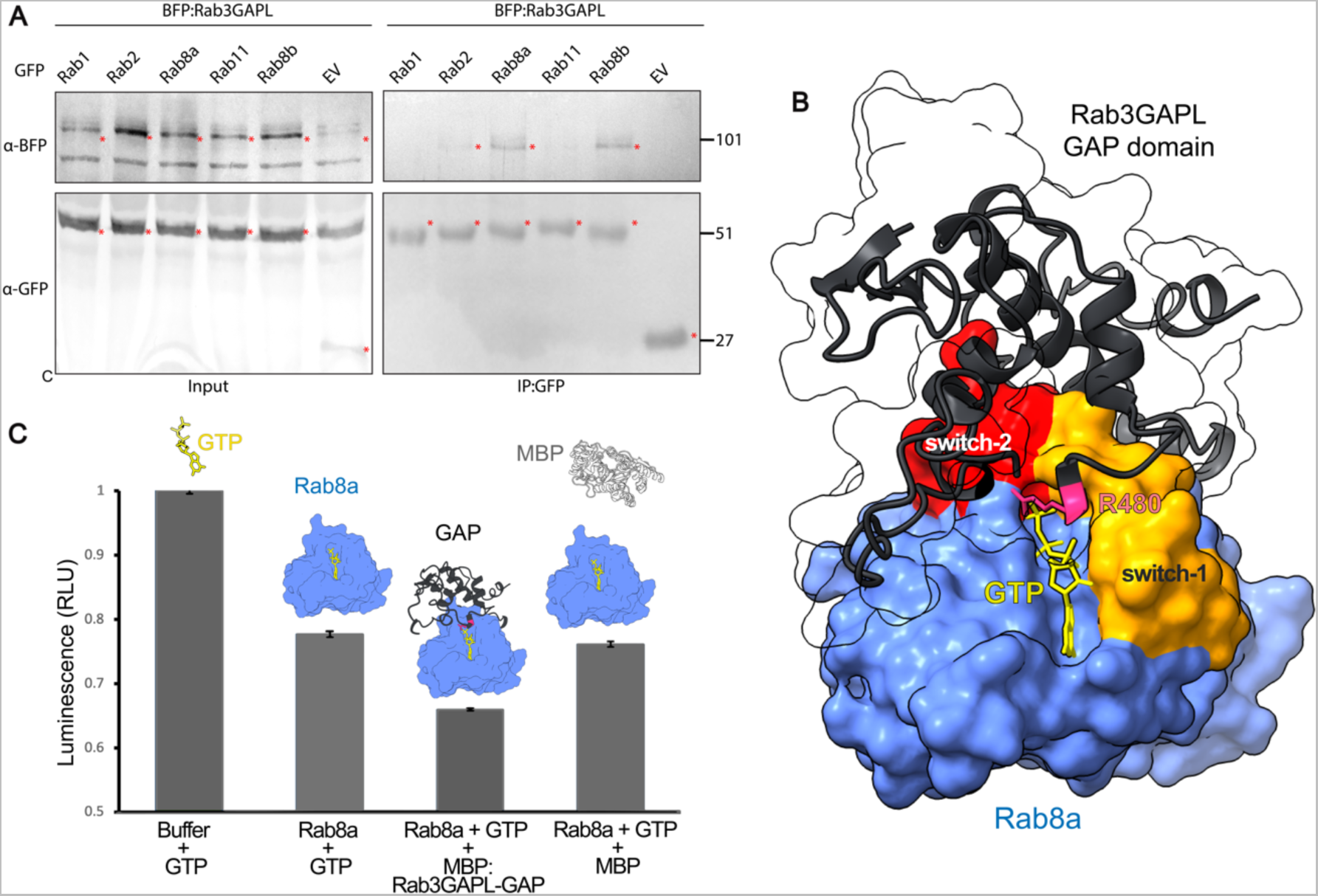
Rab3GAPL interacts with Rab8a and stimulates its GTP hydrolysis activity. (A) Rab3GAPL binds to Rab2 and Rab8 families of small GTPases *in planta*. BFP:Rab3GAPL was transiently co-expressed with either GFP:Rab1, GFP:Rab2, GFP:Rab8a, GFP:Rab11, GFP:Rab8b, or GFP:EV. IPs were obtained with anti-GFP antiserum. Total protein extracts were immunoblotted. Red asterisks indicate expected band sizes. (B) Predicted AF2-M model of Rab8a in complex with the GAP domain of Rab3GAPL. The catalytic arginine residue, R480 (magenta), of Rab3GAPL is positioned across the GTP binding pocket of Rab8a. (C) The GAP domain of Rab3GAPL stimulates GTPase activity of Rab8a. A luciferase-based GTPase assay was used to measure the amount of GTP over 120 minutes at room temperature. Bar graph shows the effect of purified MBP:Rab3GAPL GAP domain or MBP control on the GTPase activity of Rab8a across 3 repeats.

To gain further insights into the Rab3GAPL-Rab8a association, we utilized AF2-M. The predicted AF2 model suggests that the catalytic arginine (R480) of Rab3GAPL is located across the guanine nucleotide binding pocket flanking switch-1 and switch-2 regions on potato Rab8a (Fig. 4B and S4A), suggesting that Rab8a could be a substrate of Rab3GAPL in plants. Potato Rab8a displayed a high degree of protein sequence conservation, with 68% amino acid identity, and a high degree of structural similarity, with an RMSD value of 0.8, when compared to human Rab8a (Fig. S4B-C). Leveraging this structural conservation, we performed AF2-guided *ab initio* molecular replacement to obtain potato Rab8a bound to GTP. We replaced the crystal structure of human Rab8a bound to GTP (PDB:6WHE) (*41*) with the AF2 model of the potato Rab8a (Fig. S4C-D). The resulting Rab3GAPL-Rab8a-GTP model demonstrated that the catalytic arginine finger of Rab3GAPL is positioned to engage with the GTP-binding pocket of the potato Rab8a and makes contacts with the conserved glutamine from DTAGQ motif of the switch-2 region of Rab8a (Fig. 4B and S4D-E). Such interactions between the catalytic arginine and the switch-2 glutamate typically facilitate the nucleophilic attack by a water molecule on the γ-phosphate of GTP, leading to GTP hydrolysis and the subsequent release of inorganic phosphate (*42*) (Fig. S4E).

Based on these findings, we further investigated the interplay between Rab8a and Rab3GAPL by performing biochemical assays. We conducted *in vitro* GAP assays to determine whether Rab3GAPL enhances the GTP hydrolysis activity of Rab8a using proteins purified from *E. coli*. The titration of 5 μM of purified Rab8a into the GTP reaction buffer led to a significant reduction in free GTP levels compared to the buffer control (Fig. 4C), approving the functionality of purified Rab8a protein. When Rab8a was incubated together with the purified GAP domain of Rab3GAPL, we noted a more pronounced reduction in free GTP levels in the buffer compared to Rab8a alone or using maltose binding protein (MBP) control (Fig. 4C). These results indicate that Rab3GAPL promotes the GTP hydrolysis activity of Rab8a and that Rab3GAPL can function as a conventional GAP for Rab8a. In conjunction with our previous findings demonstrating the association between Rab8a and ATG8CL in autophagy activation (*15*), these results support the notion that Rab3GAPL targets Rab8a as a GAP substrate to regulate autophagy. Since Rab3GAPL suppresses autophagy in a manner that relies on its GAP and ATG8-binding activities (Fig. 2A-B, E-F), we suggest that Rab3GAPL could regulate autophagy by switching off Rab8a at autophagosome biogenesis sites, where ATG8 proteins are actively recruited (*43*).

### 5. Rab3GAPL increases susceptibility to *Phytophthora infestans* independent of its autophagy suppression activity

Given the recent findings supporting the positive role of Rab8a in autophagy and immunity against *P. infestans* (*15, 17*), we next investigated whether Rab3GAPL has any impact on pathogen resistance. Firstly, we tested if Rab3GAPL affects immunity to *P. infestans* in an autophagy-dependent manner by overexpressing the WT Rab3GAPL or its AIM mutant. In four independent experiments, infected leaf patches expressing GFP:Rab3GAPL or GFP:Rab3GAPL^AIM^ showed enhanced disease symptoms with significant increases in infection lesion sizes compared to the GFP control (Fig. 5A-B). We validated these results by performing infection assays using a red fluorescent strain of *P. infestans*, 88609td, which allows measurement of pathogen biomass through imaging of hyphal threads via fluorescent microscopy. Consistently, *P. infestans* hyphal growth was significantly higher in leaf patches overexpressing GFP:Rab3GAPL or GFP:Rab3GAPL^AIM^ compared to GFP control samples (Fig. S5A-B). Intriguingly, the overexpression of Rab3GAPL^AIM^ mutant, which is impaired in autophagy suppression and ATG8 binding, promoted infection to levels comparable to that of WT Rab3GAPL (Fig. 5A-B and S5A-B). These results indicate that Rab3GAPL negatively regulates immunity independent of its function in autophagy.

**Figure 5.**
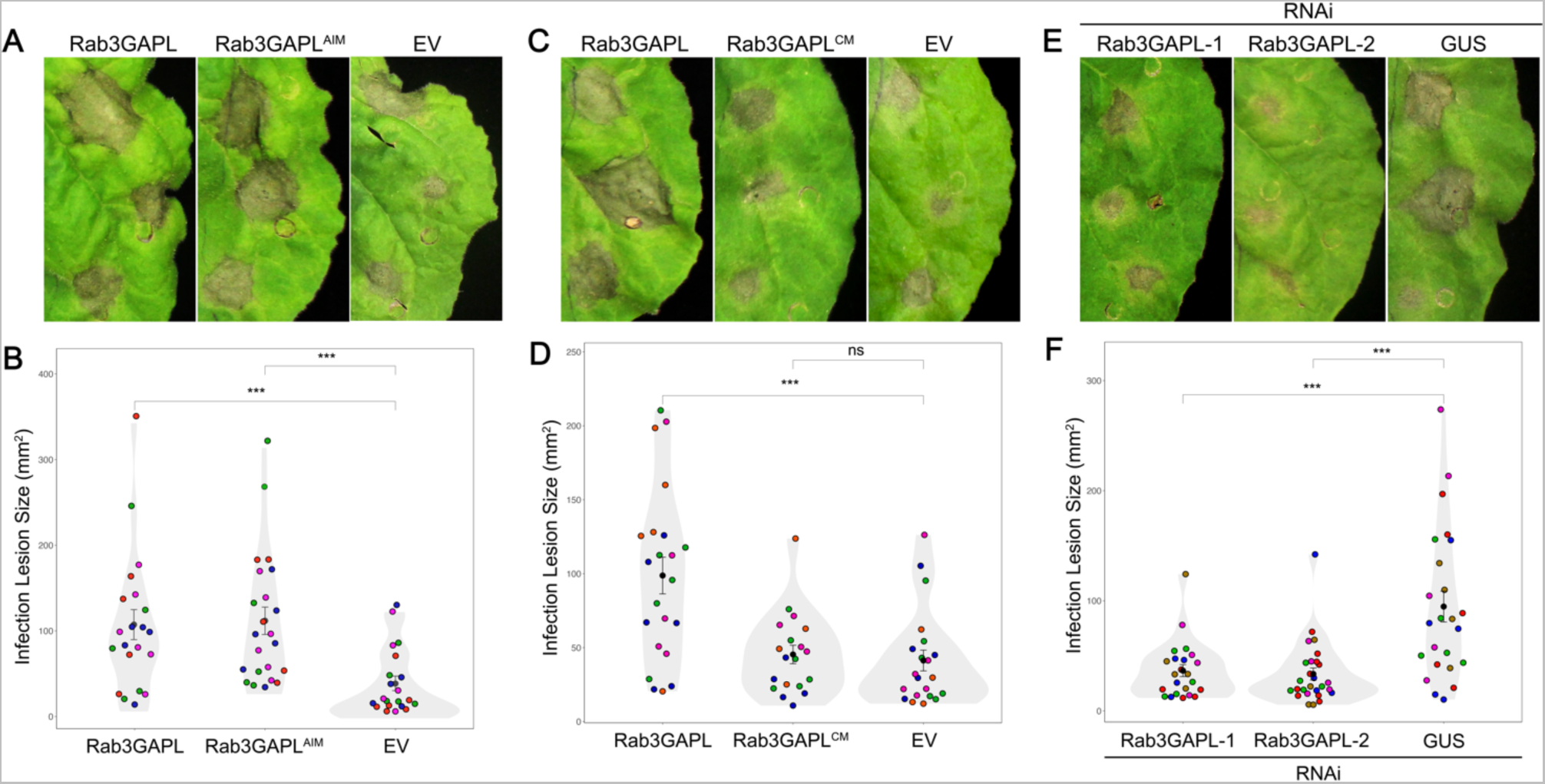
Rab3GAPL increases susceptibility to *Phytophthora infestans* in a catalytic activity-dependent, AIM-independent manner. (A-B) Rab3GAPL increases susceptibility to *P. infestans* in an AIM-independent manner. (A) *N. benthamiana* leaves expressing Rab3GAPL, Rab3GAPL^AIM^ or EV control were infected with *P. infestans*, and pathogen growth was calculated by measuring infection lesion size 7 days post-inoculation. (B) Both Rab3GAPL expression (107.3, N = 21 spots) and Rab3GAPL^AIM^ expression (111.8, N = 23 spots) significantly increase *P. infestans* lesion sizes compared to EV control (39.0, N = 21 spots). Statistical differences were analyzed by Mann-Whitney U test in R. Measurements were highly significant when p<0.001 (***). (C-D) Rab3GAPL increases susceptibility to *P. infestans* in a catalytic activity-dependent manner. (C) *N. benthamiana* leaves expressing Rab3GAPL, Rab3GAPL^CM^ or EV control were infected with *P. infestans*, and pathogen growth was calculated by measuring infection lesion size 7 days post-inoculation. (D) Rab3GAPL expression (98.8, N = 22 spots) significantly increases *P. infestans* lesion sizes compared to EV control (45.4, N = 19 spots), whereas Rab3GAPL^CM^ expression (41.3, N = 21 spots) has no significant effect compared to EV control. Statistical differences were analyzed by Mann-Whitney U test in R. Measurements were highly significant when p<0.001 (***). (E-F) Silencing *Rab3GAPL* reduces susceptibility to *P. infestans*. (E) *N. benthamiana* leaves expressing RNAi:Rab3GAPL-1, RNAi:Rab3GAPL-2 or RNAi:GUS control were infected with *P. infestans*, and pathogen growth was calculated by measuring infection lesion size 8 days post-inoculation. (F) Both RNAi:Rab3GAPL-1 expression (36.6, N = 23 spots) and RNAi:Rab3GAPL-2 expression (33.5, N = 27 spots) significantly reduce *P. infestans* lesion sizes compared to RNAi:GUS control (94.8, N = 24 spots). Statistical differences were analyzed by Mann-Whitney U test in R. Measurements were highly significant when p<0.001 (***).

Secondly, we explored whether enhanced susceptibility phenotype caused by overexpression of Rab3GAPL requires its GAP activity. Notably, overexpression of the GAP mutant (GFP:Rab3GAPL^CM^) did not cause any difference in *P. infestans* infection lesion sizes compared to the GFP control, unlike the WT GFP:Rab3GAPL construct which enhanced disease symptoms (Fig. 5C-D). These findings indicate that the enhanced pathogen growth phenotype caused by Rab3GAPL overexpression is reliant on its GAP activity, suggesting a potential negative regulation of immunity through the restriction of Rab-mediated trafficking.

Thirdly, we conducted infection assays upon downregulation of Rab3GAPL expression. To achieve this, we employed silencing constructs RNAi:Rab3GAPL-1 and RNAi:Rab3GAPL-2, designed to specifically target *Rab3GAPL* in *N. benthamiana* (Fig. S2D). In agreement with our overexpression assays, which suggested a negative role of Rab3GAPL in immunity (Fig. 5A-D), the silencing of Rab3GAPL using either RNAi:Rab3GAPL constructs significantly enhanced *P. infestans* infection lesion size and hyphal growth compared to the control construct RNAi:GUS (Fig. 5E-F and S5C-D). These results show that Rab3GAPL acts as a susceptibility factor in a catalytic activity-dependent, but AIM-independent manner. Collectively, these findings suggest that the negative regulatory function of Rab3GAPL in defense against *P. infestans* is independent of autophagy, highlighting its involvement in alternative mechanisms of immune regulation.

### 6. Rab3GAPL antagonizes Rab8a-mediated defense vesicle dynamics and secretion

Recent studies have revealed the contribution of Rab8a in defense-related secretion and basal immunity against *P. infestans*. Additionally, pathogen effectors specifically target Rab8a to undermine its immune functions, including the secretion of pathogenesis related protein-1 (PR-1) into the apoplast (*15, 44*). Given the immunosuppressive role of Rab3GAPL, which is dependent on its GAP activity but not its interaction with ATG8 (Fig. 5), we hypothesized that Rab3GAPL negatively regulates defense-related secretion mediated by Rab8a. To test this, we first assessed the impact of Rab3GAPL on defense-related secretory responses by examining its effect on PR-1 secretion. To stimulate endogenous PR-1 induction, we challenged the leaf patches expressing Rab3GAPL and controls with *P. infestans* extract, serving as a pathogen-associated molecular pattern (PAMP) cocktail. The secretion of PR-1 to the apoplast was drastically reduced in samples expressing Rab3GAPL or Rab3GAPL^AIM^ compared to the GFP vector control. However, the apoplastic levels of PR-1 were unaffected by the catalytic mutant, Rab3GAPL^CM^ (Fig. 6A). While apoplastic PR-1 levels were reduced in samples overexpressing Rab3GAPL and Rab3GAPL^AIM^, cytoplasmic PR-1 levels reciprocally increased, suggesting that the decrease in apoplastic PR-1 was not due to impaired PR-1 expression (Fig. 6A).

**Figure 6.**
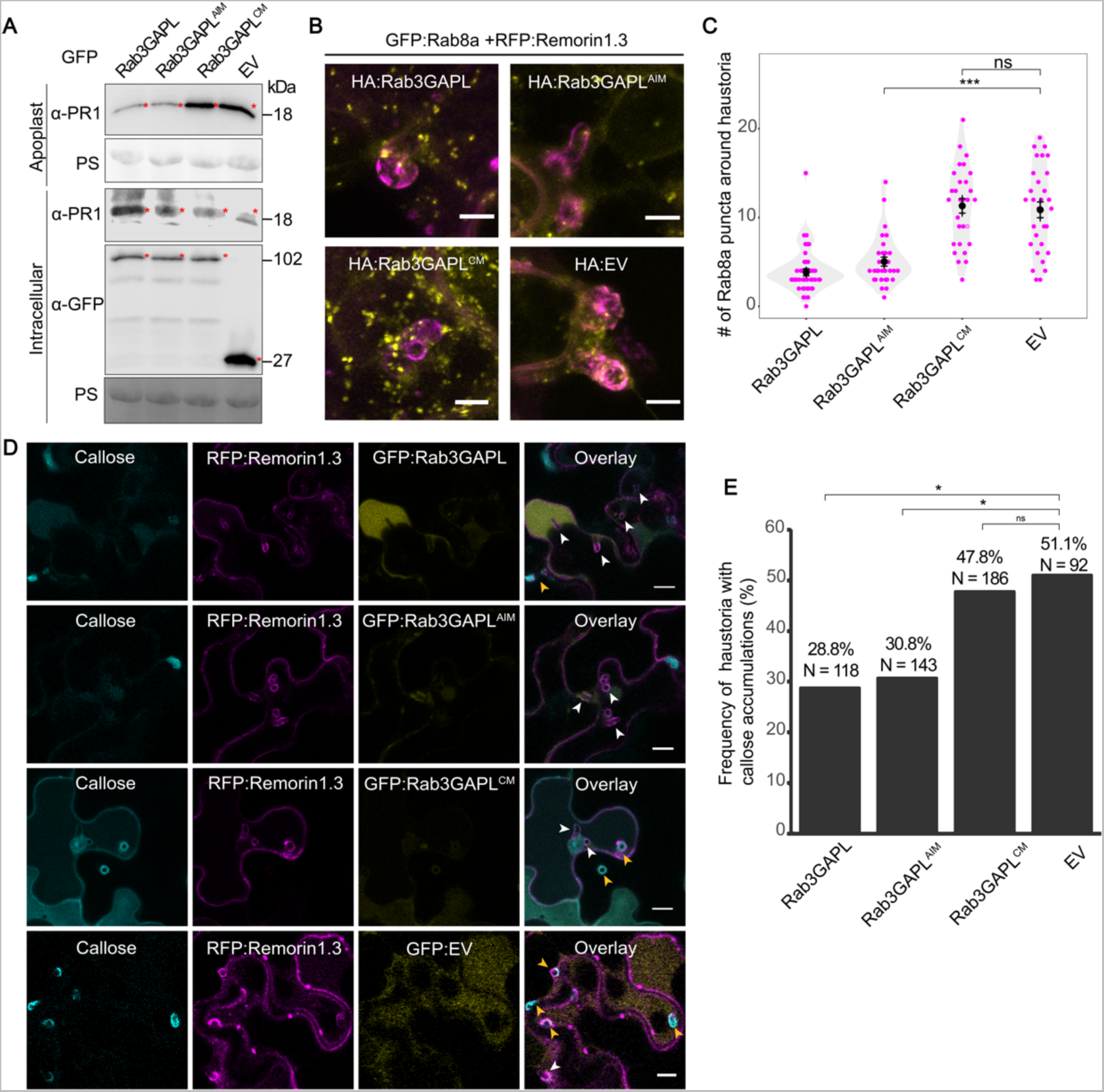
Rab3GAPL suppresses defense-related secretion in a catalytic activity-dependent manner. (A) Western blot shows Rab3GAPL and Rab3GAPL^AIM^, but not Rab3GAPL^CM^, reduces antimicrobial PR-1 secretion to the apoplast compared to EV control. *N. benthamiana* leaves were infiltrated to express GFP:Rab3GAPL, GFP:Rab3GAPL^AIM^, GFP:Rab3GAPL^CM^ or GFP:EV. The leaves were then challenged with *P. infestans* extract at 3 dpi and proteins were extracted from the apoplast and leaf tissue at 4 dpi and immunoblotted. Red asterisks show expected band sizes. (B-C) Rab3GAPL reduces the number of Rab8a vesicles around haustoria in a catalytic activity-dependent manner. (B) Confocal micrographs of *N. benthamiana* leaf epidermal cells transiently expressing GFP:Rab8a and RFP:Remorin1.3 with HA:Rab3GAPL, HA:Rab3GAPL^AIM^, HA:Rab3GAPL^CM^ or HA:EV. Images shown are maximal projections of 12 frames with 1.2 μm steps. Scale bars represent 10 μm. (C) Rab3GAPL expression (3, N = 42 haustoria) or Rab3GAPL^AIM^ expression (4, N = 31 haustoria) significantly reduce number of Rab8a vesicles around haustoria compared to EV control (11.5, N = 30 haustoria), while Rab3GAP^CM^ expression (12, N = 29 haustoria) has no significant effect compared to EV control. Statistical differences were analyzed by Mann-Whitney U test in R. Measurements were highly significant when p<0.001 (***). (D-E) Rab3GAPL reduces callose deposition at *P. infestans* haustoria in a catalytic activity-dependent manner. (D) Confocal micrographs of *N. benthamiana* leaf epidermal cells transiently expressing RFP:Remorin1.3 with GFP:Rab3GAPL, GFP:Rab3GAPL^AIM^, GFP:Rab3GAPL^CM^ or GFP:EV. The leaves were infected with *P. infestans* spores at 1 dpi, and stained with aniline blue to visualize callose at 4 dpi. Images shown are single plane images. White arrows indicate haustoria. Scale bars represent 10 μm. (E) Bar graphs showing Rab3GAPL expression (28.8%, N = 118 haustoria) or Rab3GAPL^AIM^ expression (30.8%, N = 143 haustoria) significantly reduce the frequency of callose deposition around haustoria compared to EV control (51.1%, N = 92 haustoria), while Rab3GAPL^CM^ expression (47.8%, N = 186 haustoria) has no significant effect compared to EV control. Statistical differences were analyzed by chi-squared test in R. Measurements were significant when p<0.05 (*).

Our previous work revealed that Rab8a-labeled vesicle-like structures are deposited around the extrahaustorial membrane (EHM) that envelopes the *P. infestans* haustorium (*15*). We reasoned that these Rab8a vesicles could deliver defense compounds to restrict pathogen invasion, whereas Rab3GAPL plays an antagonistic role in regulating Rab8a-mediated transport pathways. Supporting this notion, we observed a notable reduction in the abundance of Rab8a-labeled vesicles around the haustorium interface upon Rab3GAPL overexpression compared to the vector control (Fig. 6B-C). We observed a similar decrease in Rab8a vesicle abundance around haustoria when we overexpressed Rab3GAPL^AIM^. In contrast, expression of Rab3GAPL^CM^ did not affect abundance of Rab8a puncta around haustoria, behaving similarly to the vector control (Fig. 6B-C). Collectively, our findings provide evidence that Rab3GAPL negatively regulates secretory defenses dependent on Rab8a during the immune response to *P. infestans*. These results also suggest that Rab3GAPL could antagonize focal immune responses targeted to the pathogen interface.

Following up, we examined the potential impact of Rab3GAPL on plant focal immunity. Specifically, we explored whether Rab3GAPL influences callose deposition surrounding *P. infestans* haustoria. Callose deposits play a crucial role in the immune response, especially when pathogens establish specialized host-pathogen interfaces like haustoria for invading host cells (*45–47*). We observed around 40% reduction in the occurrence of haustoria with callose deposits upon Rab3GAPL overexpression compared to the vector control. We observed a similar decrease in haustoria with callose deposits when Rab3GAP^AIM^ was overexpressed, but not when Rab3GAP^CM^ was overexpressed (Fig. 7D-E). These findings, combined with our earlier observations of Rab3GAPL’s disruption of Rab8a vesicle dynamics around haustoria, suggest that Rab3GAPL negatively regulates the secretory pathways that are locally deployed at pathogen penetration sites. Moreover, these results provide insights into the potential mechanisms underlying the increased susceptibility phenotype observed towards *P. infestans* upon overexpression of Rab3GAPL and its AIM mutant, Rab3GAPL^AIM^ (Fig. 5A-B).

**Figure 7.**
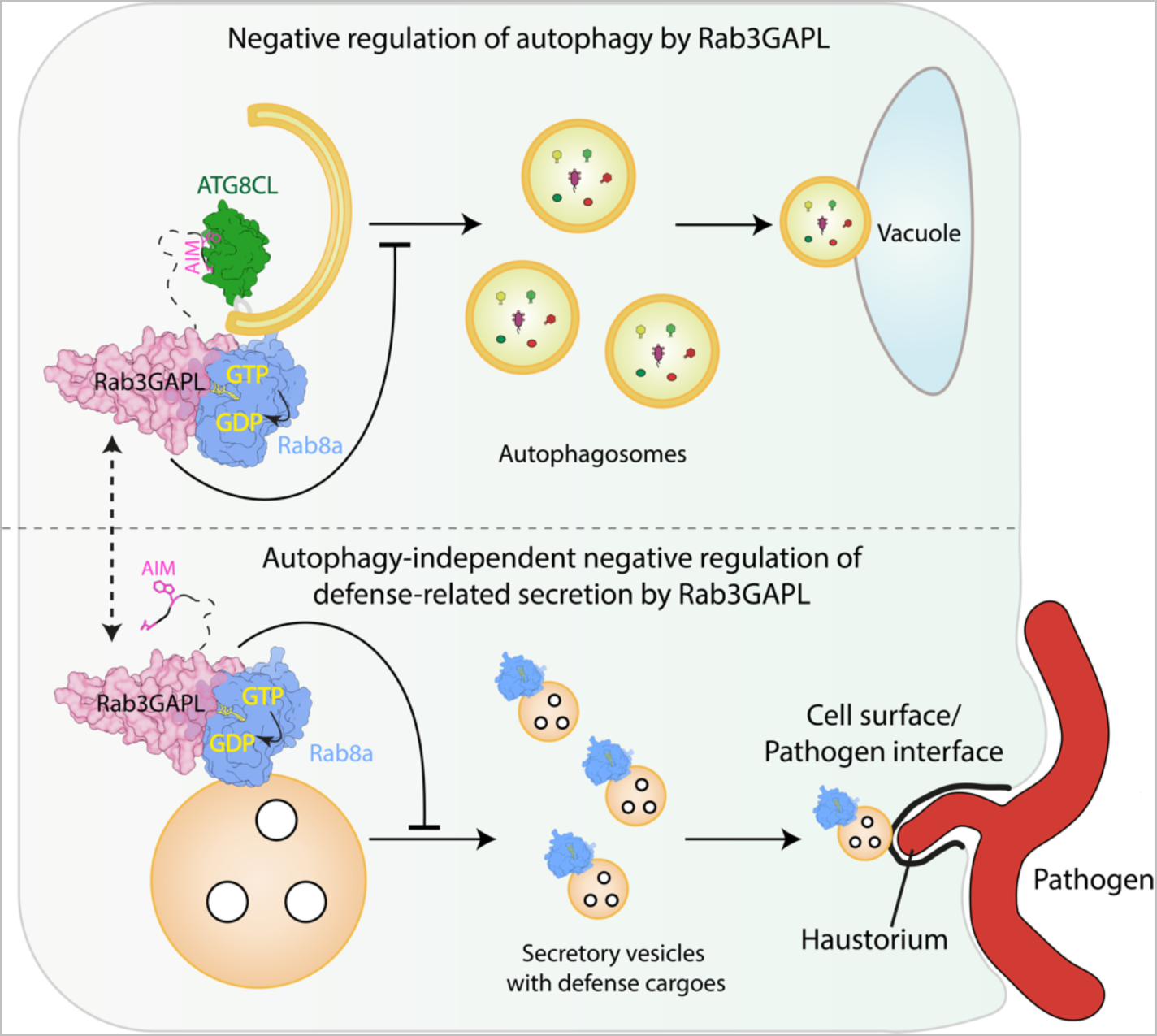
Summary model of the negative regulation of autophagy and immunity by Rab3GAPL. Rab3GAPL toggles between activities to optimize autophagy and defense-related secretion by hindering Rab8a-mediated vesicle trafficking through promoting the GTP hydrolysis of Rab8a. Rab3GAPL suppresses autophagy by binding to ATG8, the core autophagy adaptor using its ATG8-interacting motif (AIM), and deactivating Rab8a using its GTP-ase activating protein (GAP) domain. Beyond autophagy regulation, Rab3GAPL modulates Rab8a-mediated defense-related secretion towards the cell surface/pathogen interface. This immune modulation is independent of its AIM, but dependent on its GAP domain. Consequently, Rab3GAPL functions as a molecular rheostat, dynamically balancing autophagy and immune responses to maintain cellular homeostasis. This model highlights the molecular interplay between a RabGAP-Rab pair and ATG8, providing new insights into the complex membrane transport mechanisms that underpin plant autophagy and immunity.

## Discussion

In this study, we investigated the membrane trafficking processes involved in plant autophagy and immunity. Our findings revealed the role of Rab3GAPL as a regulator of vesicle transport that carries a canonical AIM to interact with ATG8 and suppress plant autophagy (Fig. 1-2 and S1-2). Although the Rab3GAPL AIM is broadly conserved in land plants, some plants in the Brassicales order exhibit mutations in their Rab3GAPL AIM residues, suggesting potential diversification in autophagy regulation (Fig. 3A-B and S3A-C). We also discovered that Rab3GAPL targets Rab8a, an important GTPase involved in autophagy activation and immunity. By stimulating Rab8a’s GTPase activity, Rab3GAPL effectively suppresses autophagy (Fig. 4).

Interestingly, our findings extend beyond autophagy regulation, as we have uncovered an additional role of Rab3GAPL in negatively modulating immunity towards *P. infestans* that is independent of its ATG8-binding activity (Fig. 5-6). This modulation relies on the GAP function of Rab3GAPL and involves the inhibition of Rab8a-mediated trafficking diverted towards the pathogen interface (Fig. 6B-E). Our results suggest a model in which Rab3GAPL impedes Rab8a-mediated vesicle trafficking by promoting Rab8a’s GTP-to-GDP switch. While Rab8a-mediated trafficking is crucial for autophagy (*15*), Rab3GAPL suppresses this process at autophagosome biogenesis sites where ATG8 is enriched. Additionally, Rab3GAPL can subvert defense-related secretion mediated by Rab8a, possibly to mitigate auto-immune responses and to adjust appropriate resource allocation (Fig.7).

Our findings uncover an intricate interplay between a RabGAP, its Rab substrate, and the core autophagy machinery, highlighting their essential roles in regulating membrane trafficking processes in both plant autophagy and immunity. The discovery of Rab3GAPL’s role in these processes, along with its potential implications in enhancing biotic and abiotic stress tolerance, presents exciting opportunities for further exploration in basic research and biotechnological applications. We envisage that continued research in this field holds promise for the development of novel strategies to fortify crop resilience, improve agricultural sustainability, and ensure global food security in the face of climate change.

## Materials and Methods

### Molecular Cloning

Molecular clonings of Rab3GAPL, Rab3GAPL^AIM^, Rab3GAPL^CM^, Rab3GAPL^CM/AIM^, Rab1, Rab2, Rab8b and Rab11 were performed using Gibson Assembly as described previously (*17, 48*). The vector backbone is a pK7WGF2 derivative domesticated for Gibson Assembly. Plasmids were constructed using primers and transformed into DH5α chemically-competent *E. coli* by heat shock. Plasmids were then amplified and extracted by PureYield™ Plasmid Miniprep System (Promega), and electroporated into *Agrobacterium tumefaciens* GV3101 electrocompetent cells. Sequencing was performed by Eurofins. RNA interference silencing constructs (RNAi:Rab3GAPL-1 and RNAi:Rab3GAPL-2) were made using an intron-containing hairpin RNA vector for RNA interference in plants (pRNAi-GG), based on Golden Gate cloning as described previously (*49*). RNAi:Rab3GAPL-1 targeted the region between 999 and 1301 bp of *Rab3GAPL*, while RNAi:Rab3GAPL-2 targeted the 3’ UTR region of *Rab3GAPL*. After amplifying the target fragments using designed primers, the fragments were inserted into the pRNAi-GG vector both in sense and anti-sense orientation using the overhangs left by BsaI cleavage. The resulting plasmid leads to expression of a construct that folds back onto itself forming the silencing hairpin structure. The subsequent steps of *E. coli* transformation, Miniprep, sequencing and agrotransformation were the same as overexpression constructs. All primers used in this study are detailed in **Table S1**. Constructs used in this study are detailed in **Table S2**.

### Plant material

Wildtype and transgenic *Nicotiana benthamiana* plants were grown in a controlled growth chamber at 24°C in a mixture of organic soil (3:1 ratio of Levington’s F2 with sand and Sinclair’s 2-5 mm vermiculite). The plants were exposed to high light intensity and subjected to a long day photoperiod consisting of 16 hours of light and 8 hours of darkness. Experiments were conducted using plants that were 4-5 weeks old.

*Marchantia polymorpha* MpEF1:MpATG8b-GFP plants expressed in Takaragaike-1 (TAK-1, male) were used. The plants were grown on half-strength Gamborg’s B5 containing 1 % agar under 50-60 mmol m^- 2^s^-1^ of white light at 22°C (*50*).

### *Phytophthora infestans* growth and infection assays

WT and tdTomato-expressing *Pytophthora infestans* 88069 isolats were grown on rye sucrose agar (RSA) media in the dark at 18°C for 10 - 15 days before harvesting zoospores (*51*). Zoospore solution was collected by adding 4°C cold water to the media and incubated at 4°C for 90 minutes. For infection assay, 10 µl droplets of zoospore solution at 50,000 spores/ml were added to the abaxial side (underside) of agroinfiltrated leaves (*52*). Leaves were then kept in humid conditions. Microscopy of infected leaves was conducted 3 days post infection. Daylight and fluorescent images were taken at 7 - 8 days post infection, which lesion sizes and hyphal growth were measured in ImageJ.

### Confocal microscopy

Confocal microscopy analyses were carried out 3 days post agroinfiltration. Leaf discs for microscopy were taken using size 4 cork borer, live-mounted on glass slides, and submerged in wells of dH2O using Carolina observation gel (Carolina Biological Supply Company). The slides were imaged using Leica TCS SP8 resonant inverted confocal microscope with 40x water immersion objective lens. The abaxial side of leaf tissue was imaged. The laser excitations for BFP, GFP and RFP tags are Diode 405 nm, Argon 488 nm and DPSS 561 nm respectively. Sequential scanning between lines was performed to prevent spectral mixing from different fluorophores when imaging samples with more than one tag. Confocal images, including Z-stack and single plane images, were analysed in ImageJ.

### Structural and sequence analyses

AF2-multimer was utilized through a subscription to the Google Colab in accordance with their guidelines (*53*). The align command in UCSF Chimera (version 1.17) was employed to superimpose the AF2 predictions onto known structures and to show the confidence score of the AF2 predictions using the local distance difference test (pLDDT) scores on the lDDT-Cα metric (*54*). The scoring scale ranges from 0 to 100, where 100 corresponds to the highest confidence values. Sequence alignment was performed using the MUSCLE algorithm (*55*), and the resulting alignments were visualized and color- coded via ESPript 3.0 (*39*) The proteins and sequences used for AF2 are detailed in **Table S3**.

### Agrobacterium-mediated transient gene expression in *N. benthamiana*

Agrobacterium-mediated transient gene expression was conducted using agroinfiltration as previously described (*56*). *Agrobacterium tumefaciens* containing the desired plasmid was washed in water and resuspended in agroinfiltration buffer (10 mM MES, 10 mM MgCl_2_, pH 5.7). BioPhotometer spectrophotometer (Eppendorf) was used to measure the OD_600_ of the bacterial suspension. This suspension was adjusted to a desired OD_600_ depending on the construct and the experiment, and then infiltrated into 3 to 4-week-old *N. benthamiana* leaf tissue using needleless 1ml Plastipak syringe.

### RNA isolation, cDNA synthesis, and RT-PCR

For RNA extraction, 56 mg of leaf tissue was frozen in liquid nitrogen. RNA was extracted using TRIzol RNA Isolation Reagent (Invitrogen) according to the user manual. RNA concentration was measured using NanoDrop Lite Spectrophotometer (Thermo Scientific). 2 μg of extracted RNA was treated with RQ1 RNase-Free DNAse (Promega), then used for cDNA synthesis using SuperScript IV Reverse Transcriptase (Invitrogen). cDNA was amplified using Phusion High-Fidelity DNA Polymerase (New England Biolabs). GAPDH level was used as a transcriptional control.

### Callose staining

Callose staining was performed as described previously (*56*). *N. benthamiana* leaf discs from infected tissue expressing the proteins of interest were collected and rinsed twice in 50% ethanol. They were then rinsed in a sodium phosphate buffer (0.07 M, pH 9.0) for 30 minutes at room temperature. The leaf discs were then incubated with 0.05% w/v aniline blue solution in the phosphate buffer for 60 minutes in the dark at room temperature. Confocal microscopy was followed afterwards.

### Co-immunoprecipitation and immunoblot analyses

Proteins were transiently expressed by agroinfiltration in *N. benthamiana* leaves and harvested 3 days post agroinfiltration. For western blotting experiments, 6 leaf discs were excised using size 4 cork borer (42 mg). For co-immunoprecipitation experiments, 2 g of leaf tissues were used. Protein extraction, purification and immunoblot analysis were performed as described previously (*17, 56*). Monoclonal anti-GFP produced in rat (Chromotek), polyclonal anti-GFP produced in rabbit (Chromotek), monoclonal anti-RFP produced in mouse (Chromotek), polyclonal anti-tBFP produced in rabbit (Evrogen) were used as primary antibodies. Anti-mouse antibody (Sigma-Aldrich), anti-rabbit antibody (Sigma-Adrich), anti-rat antibody (Sigma-Aldrich) were used as secondary antibodies. Information of antibodies is detailed in **Table S4**.

### Apoplast extraction

Apoplastic proteins were extracted as described previously (*57*). Infiltrated *N. benthamiana* leaves were detached and washed in distilled water, then rolled up and inserted in a needleless syringe filled with distilled water. The whole leaves were then infiltrated with the water by creating a negative pressure environment inside the syringe. The leaves were then centrifuged for 10 minutes at 1000g in a Falcon tube. Apoplastic washing fluid was collected at the bottom of the tube and snap-frozen in liquid nitrogen. Leaf tissue was collected from the remaining leaf for immunoblotting analysis.

### *Arabidopsis thaliana* carbon starvation assays and protein extraction

Arabidopsis seedlings of the indicated genotypes were surface sterilized and added to 3 ml of liquid ½ MS + 1% sucrose, stratified for 24 hours, and grown under constant light with gentle shaking for six days. Seedlings were then washed twice in 3 ml of ½ MS medium with or without 1% sucrose, then left to grow in either ½ MS + 1% sucrose under constant light or in ½ MS without sucrose wrapped in aluminum foil for 24 hours. Proteins were extracted in 2x Laemmli buffer (100mM Tris-HCl pH 6.8, 4% SDS, 20% glycerol, 0.01% bromophenol blue, 1.5% β-mercaptoethanol), treated at 95°C for 5 minutes and quantified using amido black precipitation. For each replicate, 20 µg of total protein were loaded per sample.

### *Marchantia polymorpha* heat stress assays and protein extraction

Two-week old plants on Gamborg’s B5 medium containing 1% agar were transformed to 37°C room for 6 hours. After 2 hours recovery, samples were collected in liquid nitrogen. GTEN buffer (10% glycerol, 50 mM Tris/HCl pH7.5, 1 mM EDTA, 300 mM NaCl, 1 mM DTT, 0.1% [v/v] Nonidet P-40/Igepal, Roche cOmplete^TM^ protease inhibitor) was added to grinded-frozen samples. After vortex, samples were centrifuged at max speed for 15 minutes at 4°C for clearing the lysates. Proteins were extracted in 4x Laemmli Buffer (116 mM Tris-HCl pH 6.8, 8% SDS, 4.9% glycerol, 0.01% bromophenol blue, 10 mM DTT) and denatured at 95°C for 5 minutes. Protein concentration was equally adjusted using amido black precipitation. 10 µg of total protein were loaded per sample.

### ATG8CL expression and purification

DNA encoding ATG8CL was amplified from GFP:ATG8CL and cloned into the vector pOPINF, generating a cleavable N-terminal 6xHis-tag with ATG8CL. Recombinant proteins were produced using *E. coli* strain BL21 (DE3) grown in lysogeny broth at 37°C to an OD_600_ of 0.6 followed by induction with 1 mM IPTG and overnight incubation at 18°C. Pelleted cells were resuspended in buffer A (50 mM Tris-HCl pH 8, 500 mM NaCl and 20 mM imidazole) and lysed by sonication. The clarified cell lysate was applied to a Ni2+- NTA column connected to an AKTA Xpress system. ATG8CL was step-eluted with elution buffer (buffer A containing 500 mM imidazole) and directly injected onto a Superdex 75 26/600 gel filtration column pre-equilibrated in buffer C (20 mM HEPES pH 7.5, 150 mM NaCl). The fractions containing ATG8CL were pooled and concentrated (concentration determined using a calculated molar extinction coefficient of 7680 M^-1^cm^-1^ for ATG8CL).

### Rab8a and Rab3GAPL-GAP expression and purification

Recombinant proteins were produced using *E. coli* strain Rosetta2 (DE3) pLysS grown in 2xTY media at 37°C to an OD600 of 0.4 – 0.6 followed by induction with 300 μM IPTG and overnight incubation at 18°C. Pelleted cells were resuspended in lysis buffer (100 mM sodium phosphate pH 7.2, 300 mM NaCl, 1 mM DTT) containing Roche cOmplete^TM^ protease inhibitor and sonicated. The clarified lysate was first purified by affinity, by using HisTrap FF (GE HealthCare) columns. The proteins were eluted with a lysis buffer containing 250 mM Imidazole. The proteins were separated by Size Exclusion Chromatography with HiLoad® 16/600 Superdex 200 pg or HiLoad 16/600 Superdex 75 pg, which were previously equilibrated in 50 mM sodium phosphate pH 7.0, 100 mM NaCl. The proteins were concentrated using Vivaspin concentrators (10000 or 30000 MWCO). Protein concentration was calculated from the UV absorption at 280 nm by DS-11 FX+ Spectrophotometer (DeNovix).

### GST pull-down assays

Pulldown experiments were performed with *E. coli* lysates as previously described (*58*). Briefly, recombinant proteins were produced using *E. coli* strain Rosetta™ 2(DE3) pLysS grown in 2x TY media at 37°C to an OD_600_ of 0.4 – 0.6 followed by induction with 300 μM IPTG and overnight incubation at room temperature. Pelleted cells were resuspended in lysis buffer (100 mM sodium phosphate pH 7.2, 300 mM NaCl, 1 mM DTT) containing Roche cOmplete^TM^ protease inhibitor and sonicated. 5 μl of glutathione magnetic agarose beads (Pierce Glutathione Magnetic Agarose Beads, Thermo Fisher) were equilibrated with wash buffer (100 mM sodium phosphate pH 7.2, 300 mM NaCl, 1 mM DTT, 0.01% (v/v) IGEPAL). Clarified *E. coli* lysates were mixed with the washed beads and incubated on an end-over-end rotator for 1 hour at 4°C. Beads were washed five times with 1 ml wash buffer. Bound proteins were eluted by adding 50 μl Laemmli buffer. Samples were analyzed by immunoblotting analyses.

### Isothermal titration calorimetry (ITC)

Calorimetry experiments were carried out at 15°C in 20 mM HEPES pH 7.5, 500 mM NaCl, using an iTC200 instrument (MicroCal Inc.). For protein:peptide interactions, the calorimetric cell was filled with 90 μM ATG8CL and titrated with 1 mM Rab3GAPL-AIMp (TPVENDWTIV) or Rab3GAPL-mAIMp peptide (TPVENDATIA) from the syringe. A single injection of 0.5 μl of peptide was followed by 19 injections of 2 μl each. Injections were made at 150 seconds intervals with a stirring speed of 750 rpm. For the heats of dilution control experiments, equivalent volumes of Rab3GAPL-AIMp or Rab3GAPL-mAIMp peptide were injected into the buffer using the parameters above. The titrations were performed at 25°C, but otherwise as above. The raw titration data were integrated and fitted to a one-site binding model using the MicroCal Origin software.

### GTPase activity assay

To analyze the effect of Rab3GAPL on the GTPase activity of Rab8a, we used a luciferase-based GTPase assay (GTPase-GloTM Assay Kit by Promega). The assay was carried out as per manufacturer’s instructions. 12.5 μl of 2X GTP-GAP solution was prepared containing 5 μM GTP and 1 mM DTT in GTPase/GAP buffer. The solution was mixed with purified MBP:Rab3GAPL GAP domain. 12.5 μl of 5 μM Rab8a was added to each well. The GTPase reaction was initiated by adding 12.5 μl of the 2X GTP-GAP solution to each well. The reaction was incubated for 120 minutes at RT with shaking. 25 μl of reconstituted GTPase-Glo^TM^ Reagent was added to the completed GTPase reaction, which the remaining GTP was converted to ATP. Plate was incubated for 30 minutes at RT with shaking. Then, 50 μl of Detection Reagent was added to all the wells, and incubated for 10 minutes at RT. Finally, luminescence was measured using BioTek Synergy4 plate reader.

### CRISPR/Cas9 construct design in *Marchantia polymorpha*

Two sgRNAs were designed based on target sequence (Fig. S3E). Both sgRNAs were cloned into pMpGE_04 entry vector flanked by attL1 and attL2 sequences (*59*). Transformants were sequenced and inserted into the pMpGE010 destination vector by LR Clonase II Enzyme Mix. This vector was incorporated into *A. tumefaciens* GV3101+pSoup, which was used to transform GFP-ATG8b-TAK1. Transformants were selected on 10 µM hygromycin (*50*), genotyped and sequenced to verify mutations.

### Image processing and data analysis

Confocal microscopy images were processed with Leica LAS AF software and ImageJ. Confocal images can be single plane images or Z-stack images depending on the experiment, which is detailed in the figure legends. To quantify autophagosome punctate structures in one channel, the Z stacks were separated into individual images using ImageJ and analyzed. The counting procedure was based on the maxima function in ImageJ to avoid cytoplasmic noises. Violin plots were generated using R, bar graphs were generated using Microsoft Excel. Statistical differences were conducted in R using Student’s t-test, Welch’s t-test, Mann–Whitney U test or chi-squared test depending on the experiment, based on statistical normality and variance. Measurements were significant when p < 0.05 (*), p < 0.01 (**) and highly significant when p < 0.001 (***). All statistical calculations are detailed in **Table S5**.

### Accession numbers

Rab3GAPL (Nbe.v1.s00030g03060); StRab1(RabD2a) (PGSC0003DMP400023158); StRab2 (RABB1b) (PGSC0003DMP400022392); NbRab8b (cloned from cDNA, similar to Niben101Scf02606g00015.1); NbRab11 (Nbe.v1.s00040g37530)

## Supplementary Information

**Figure S1A.**
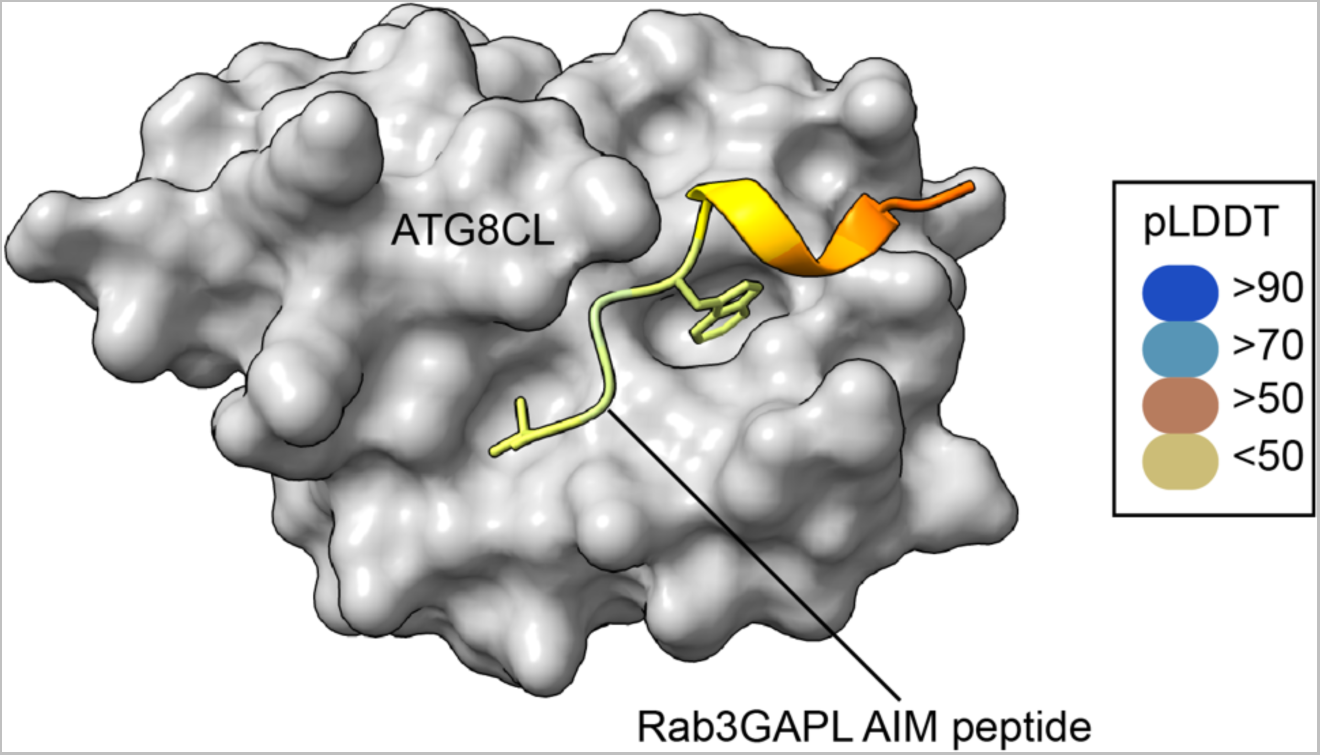
AF2-M predicted structural model of Rab3GAPL and ATG8CL interaction displaying the docking of the AIM peptide of Rab3GAPL to the AIM pocket of ATG8CL. The colors of Rab3GAPL AIM peptide are based on the AF2-calculated prediction confidence score (pLDDT) as indicated in the rectangular box.

**Figure S1B.**
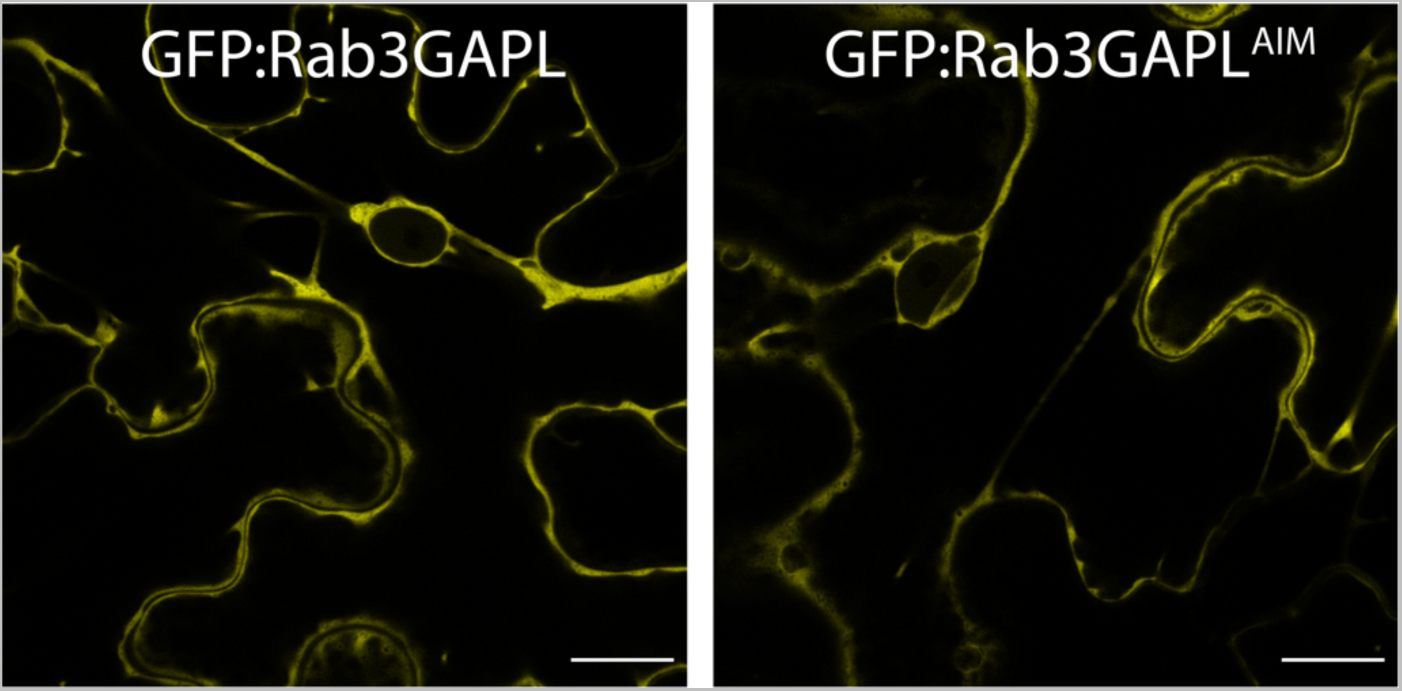
Rab3GAPL and Rab3GAPL^AIM^ show nucleus-excluded cytoplasmic localisation. Confocal micrographs of *N. benthamiana* leaf epidermal cells transiently expressing GFP:Rab3GAPL or GFP:Rab3GAPL^AIM^. Images shown are single plane images. Scale bars, 10 μm.

**Figure S2A.**
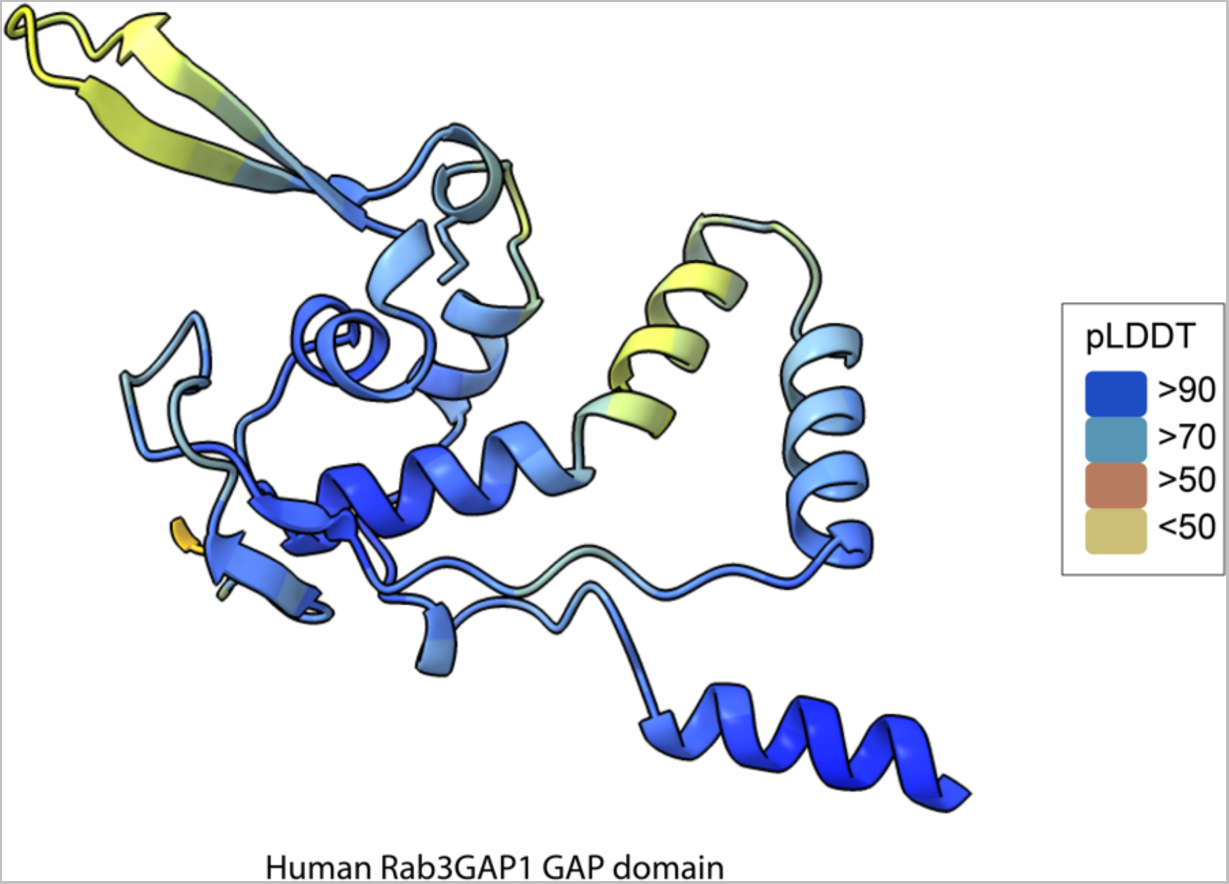
AF2 structure of the GAP domain of human Rab3GAP1. The colors of the human Rab3GAP1 GAP domain are based on the AF2-calculated prediction confidence score (pLDDT) as indicated in the rectangular box.

**Figure S2B.**
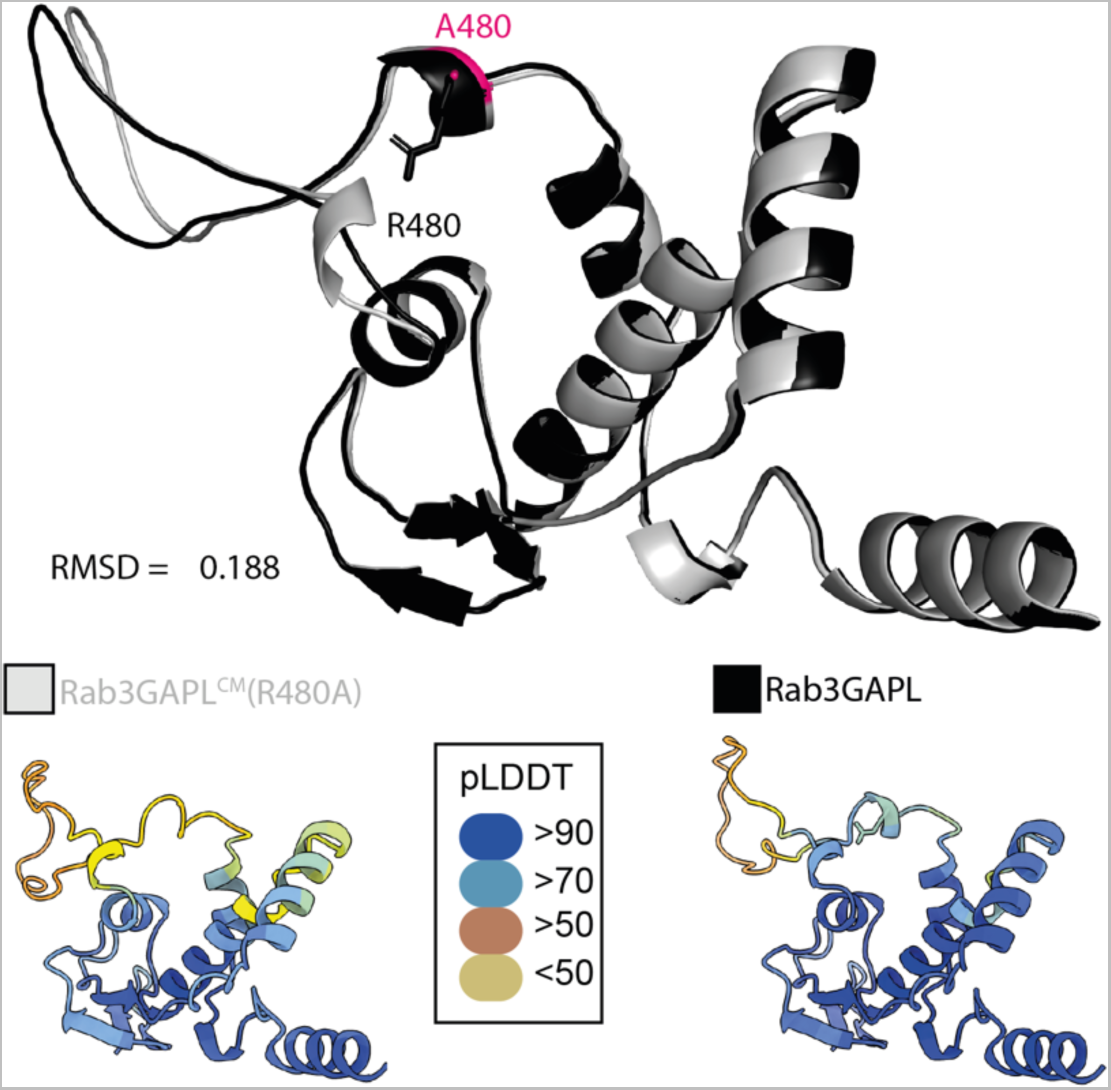
Structural alignment of Rab3GAPL and its catalytic mutant Rab3GAPLCM. Structural predictions were obtained via AF2. Model shows conservation of the overall protein structure. The colors of Rab3GAPL and Rab3GAPLCM are based on the AF2-calculated prediction confidence score (pLDDT) as indicated in the rectangular box.

**Figure S2C.**
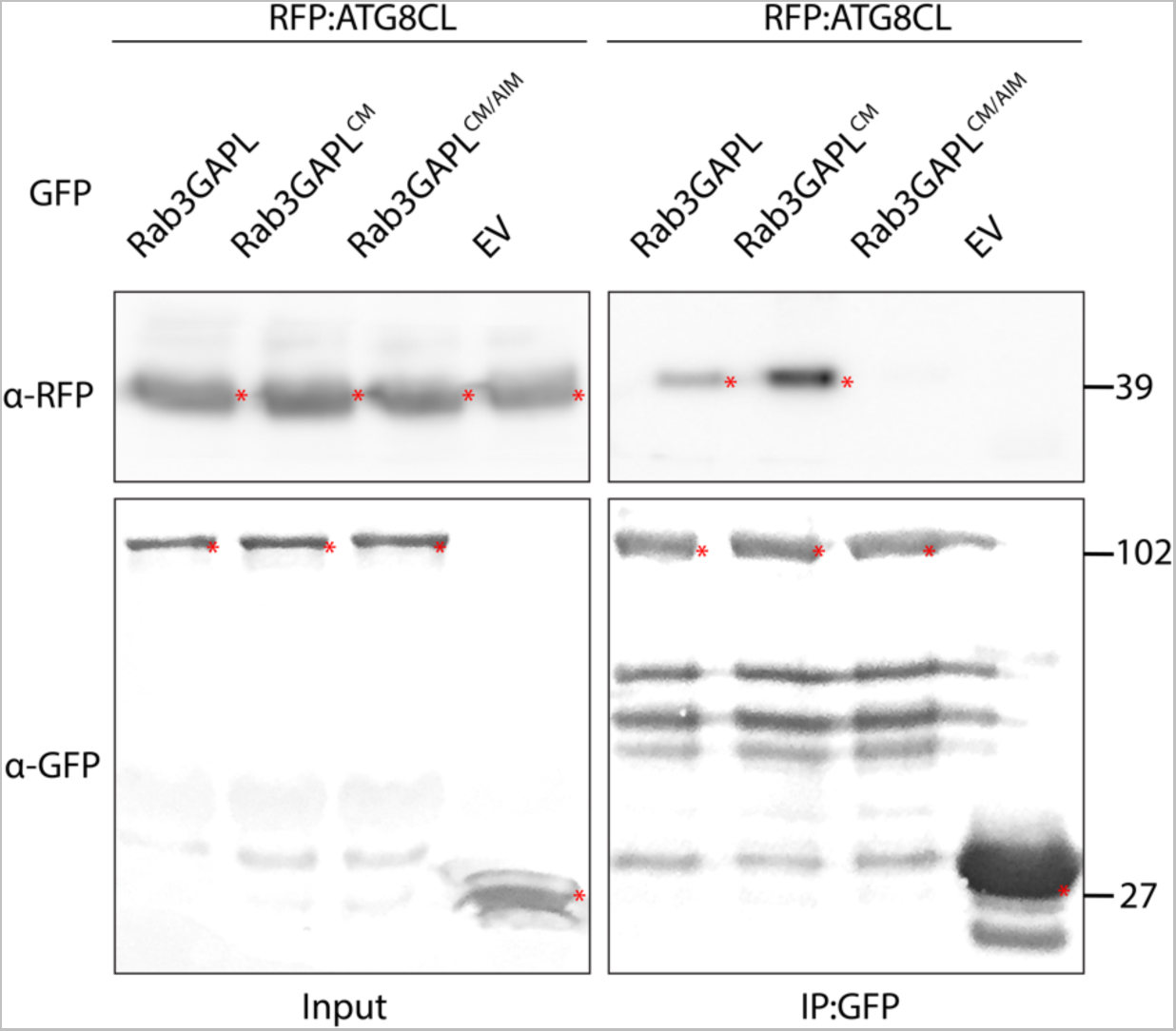
Rab3GAPL and Rab3GAPL^CM^ bind to ATG8CL via its AIM in planta. RFP:ATG8CL was transiently co-expressed with either GFP:Rab3GAPL, GFP:Rab3GAPL^CM^, GFP:Rab3GAPL^CM/AIM^ or GFP:EV. IPs were obtained with anti-GFP antiserum. Total protein extracts were immunoblotted. Red asterisks indicate expected band sizes.

**Figure S2D.**
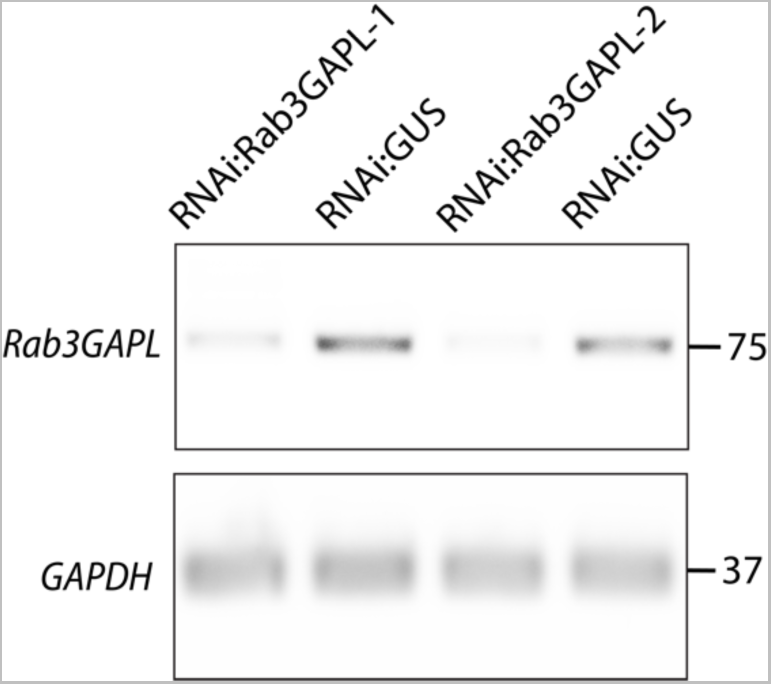
Validation of *Rab3GAPL* silencing by RNAi:Rab3GAPL-1 and RNAi:Rab3GAPL-2. Constructs carrying hairpin plasmids (pRNAi-GG) targeting Rab3GAPL-1, Rab3GAPL-2 or GUS reporter gene were infiltrated to *N. benthamiana* leaves. The expression of targeted genes was assessed by RT-PCR at 4 days post agroinfiltration. RT-PCR confirmed efficient gene silencing of *Rab3GAPL* using both RNAi:Rab3GAPL-1 and RNAi:Rab3GAPL-2 constructs. Glyceraldehyde 3-phosphate dehydrogenase (GAPDH) was used as an internal control. cDNA was synthesized using total RNA.

**Figure S2E.**
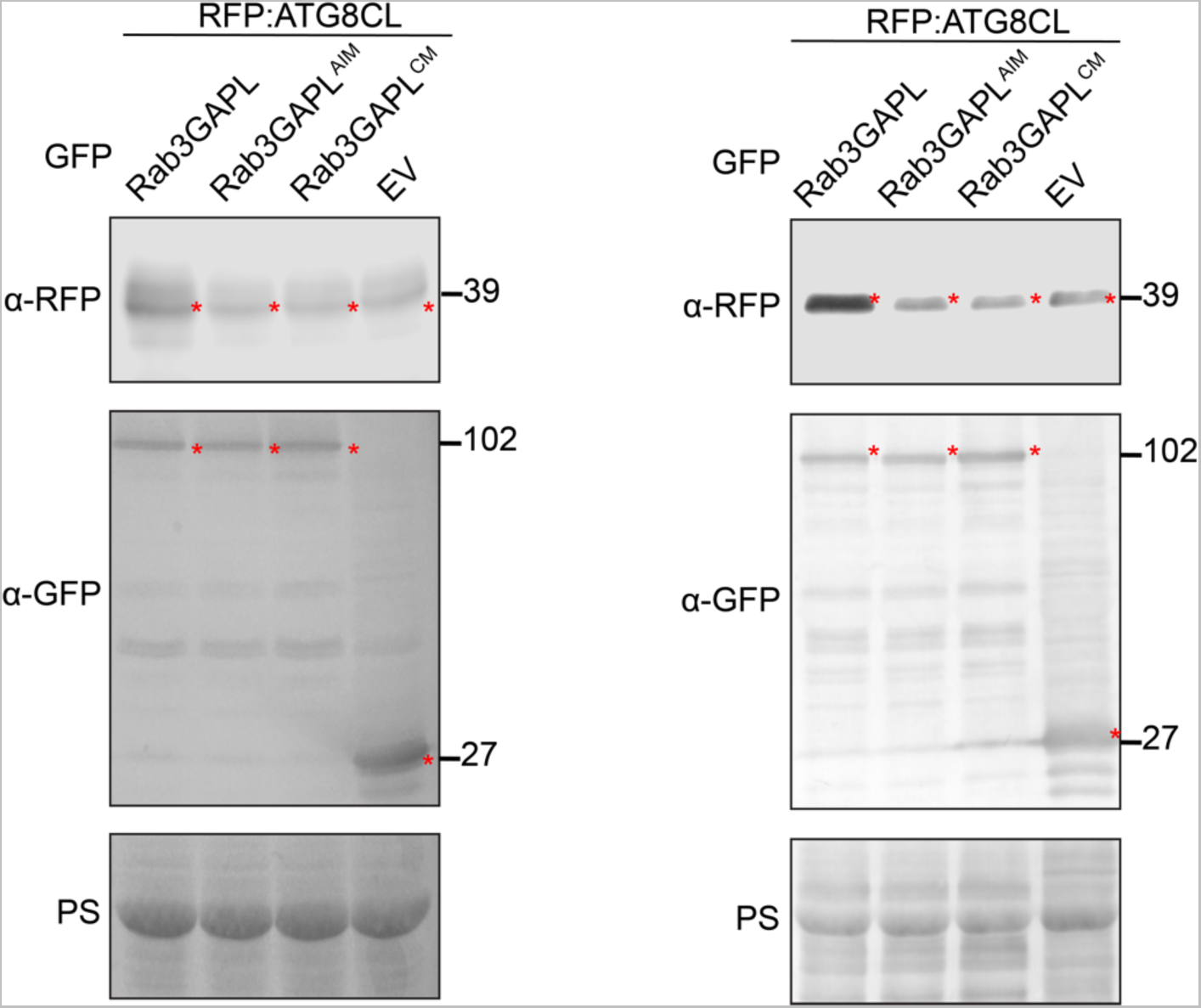
Western blots showing depletion of RFP:ATG8CL is reduced by GFP:Rab3GAPL compared to GFP:Rab3GAPL^AIM^, GFP:Rab3GAPL^CM^, or EV control (additional repeats for Figure 2I). Total protein extracts were prepared 4 days post agroinfiltration and immunoblotted. Red asterisks show expected band sizes.

**Figure S2F.**
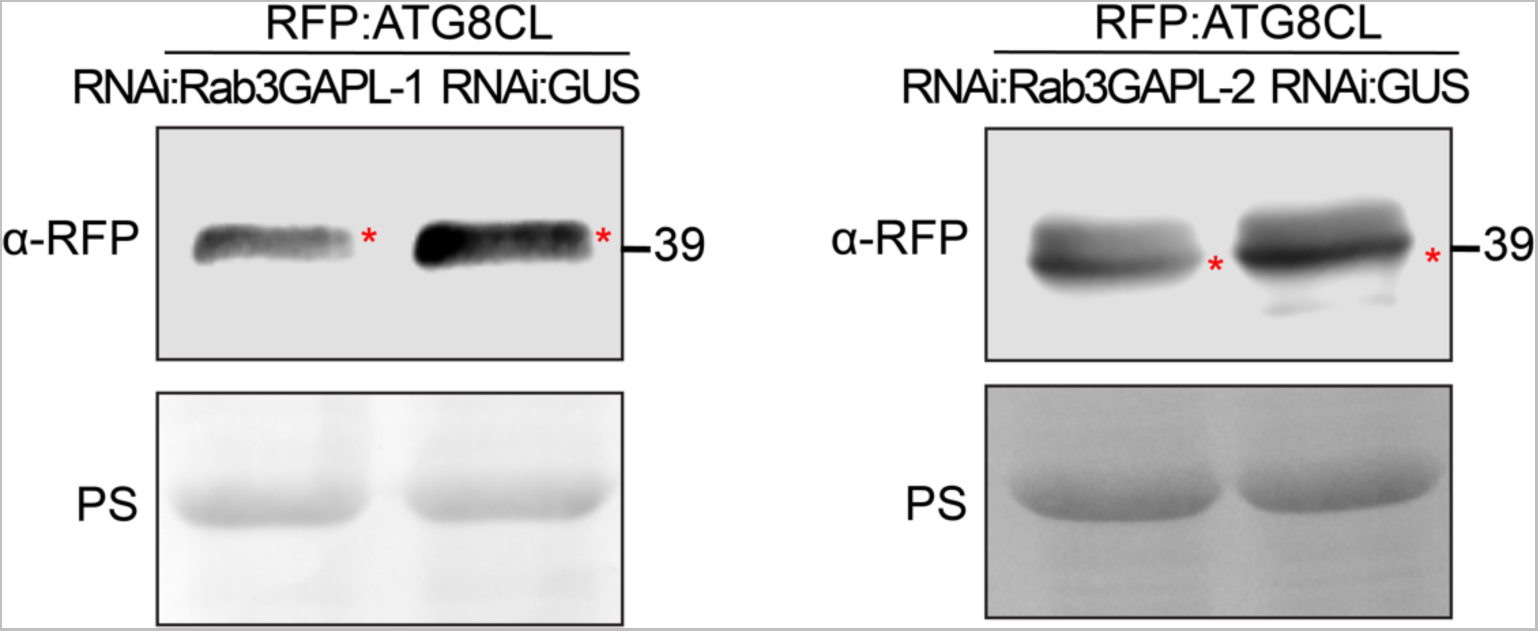
Western blots showing depletion of RFP:ATG8CL is increased by silencing Rab3GAPL-1 and -2 compared to GUS silencing control (additional repeats for Figure 2J and 2K). Total protein extracts were prepared 4 days post agroinfiltration and immunoblotted. Red asterisks show expected band sizes.

**Figure S3A.**
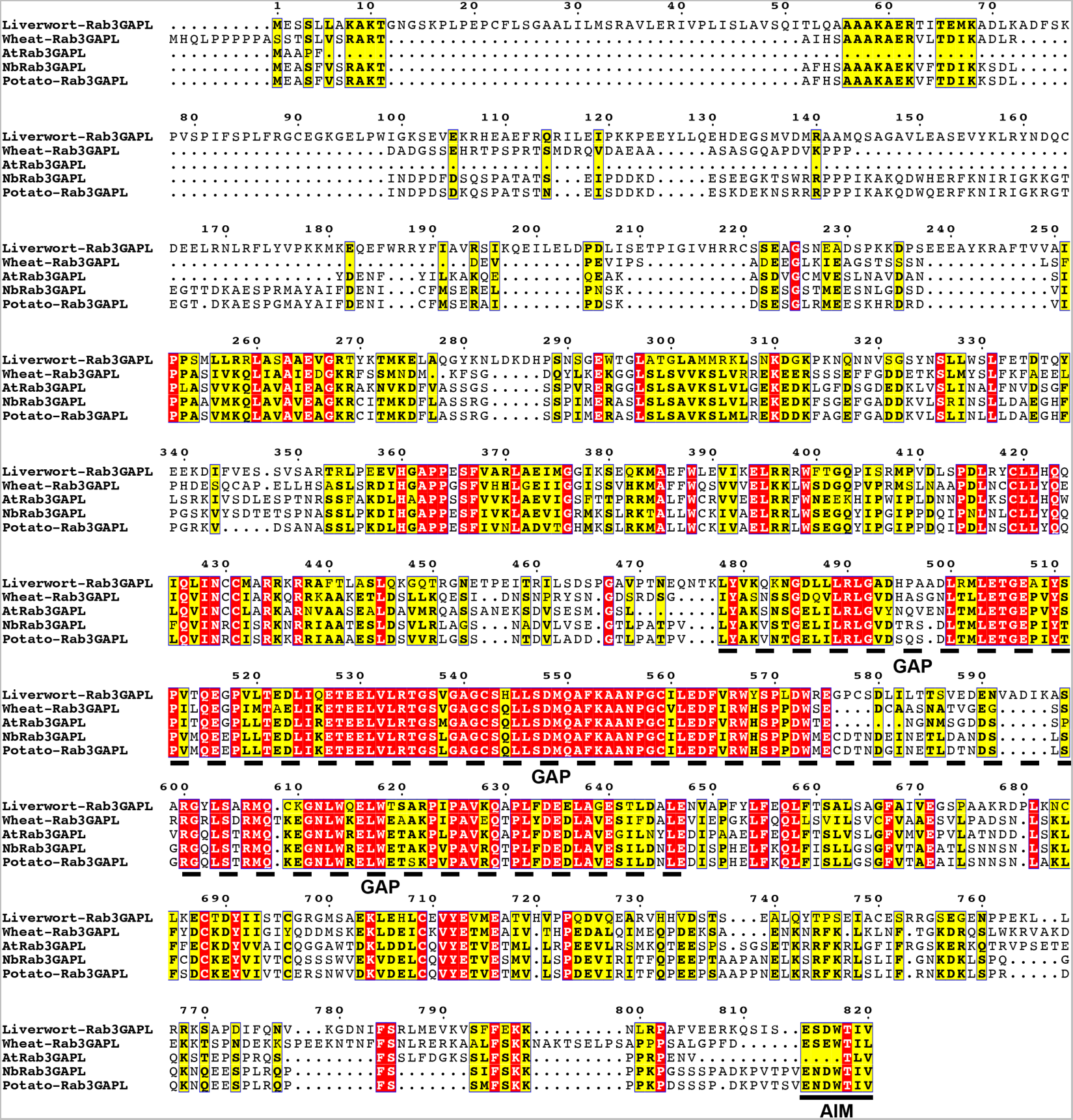
Pairwise sequence alignment comparisons of Rab3GAPLs in wheat, Arabidopsis, *N. benthamiana*, potato, and the liverwort *Marchantia polymorpha*. Alignments were obtained using the MUSCLE algorithm and were visualized and color-coded via ESPript 3.0 (*39*). The GAP domain and the AIM are illustrated using dotted and straight lines, respectively.

**Figure S3B.**
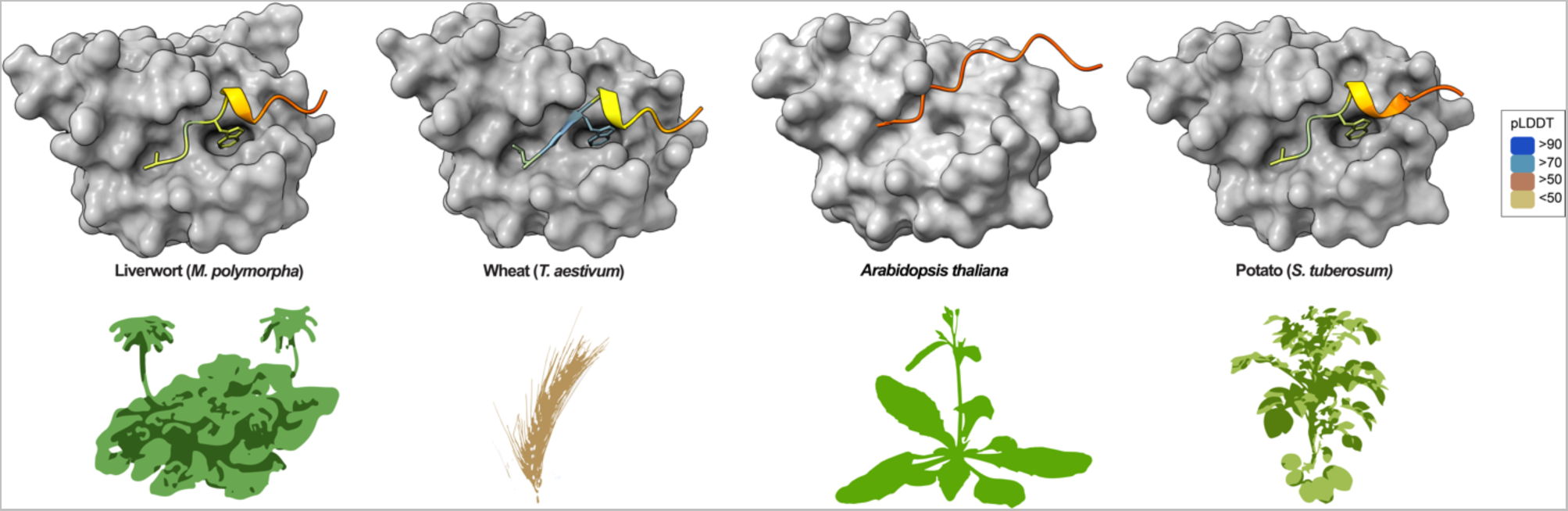
AF2-M predictions of ATG8CL with Rab3GAPL AIM sequences from liverwort (*M. polymorpha*), wheat (*Triticum aestivum*), *A. thaliana* and potato (*S. tuberosum*). Predicted models suggest AIM docking sites (W and L pockets, colored yellow and blue, respectively) on ATG8CL are associated with all tested AIMs except for the Arabidopsis Rab3GAPL AIM sequence. The colors of the AIM sequences are based on the AF2-calculated prediction confidence score (pLDDT) as indicated in the rectangular box.

**Figure S3C.**
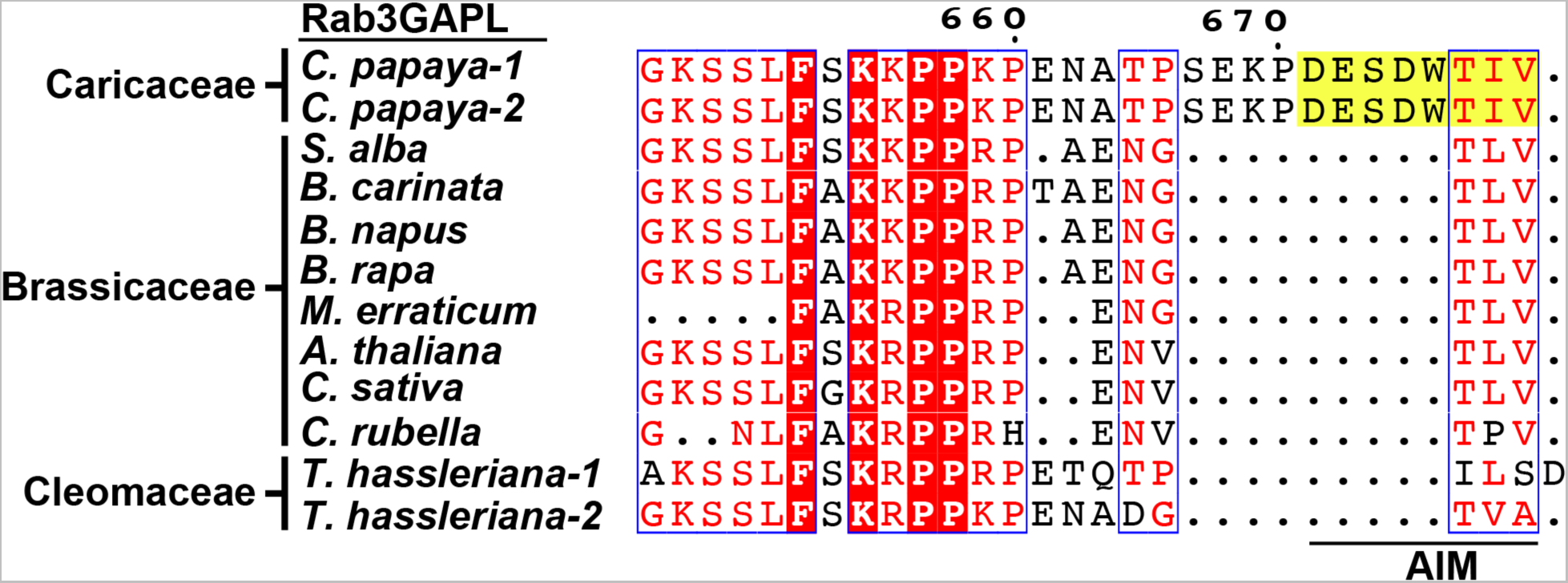
Pairwise sequence alignment comparisons of the AIMs of Rab3GAPLs in the Brassicales order of flowering plants, including the families Caricaceae, Brassicaceae and Cleomaceae. While plants in Caricaceae carry an intact AIM, plants in Brassicaceae and Cleomaceae had deletions in their AIM residues. Alignments were obtained using the MUSCLE algorithm and were visualized and color-coded via ESPript 3.0 (*39*).

**Figure S3D.**
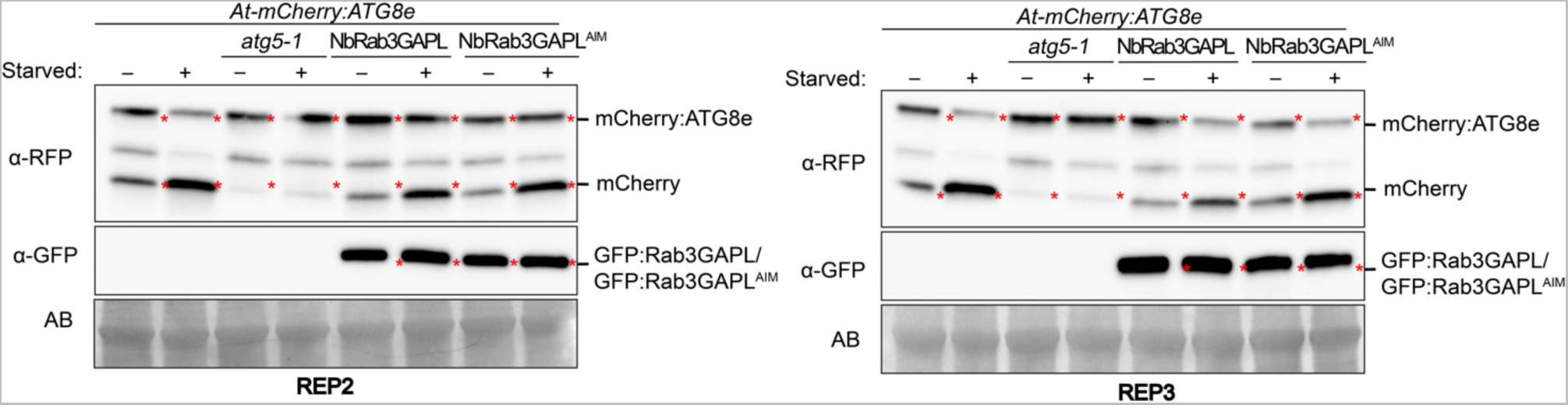
*Arabidopsis thaliana* Rab3GAPL expression mutants have reduced ATG8 autophagic flux (additional repeats for Figure 3C). Autophagic flux is measured as the ratio between free mCherry to full size mCherry:ATG8e. GFP:Rab3GAPL expression leads to reduced mCherry/mCherry:ATG8e protein signal ratio in both carbon starvation and control conditions compared to control plants. Protein extracts were prepared using 6-day-old seedlings and immunoblotted.

**Figure S3E.**
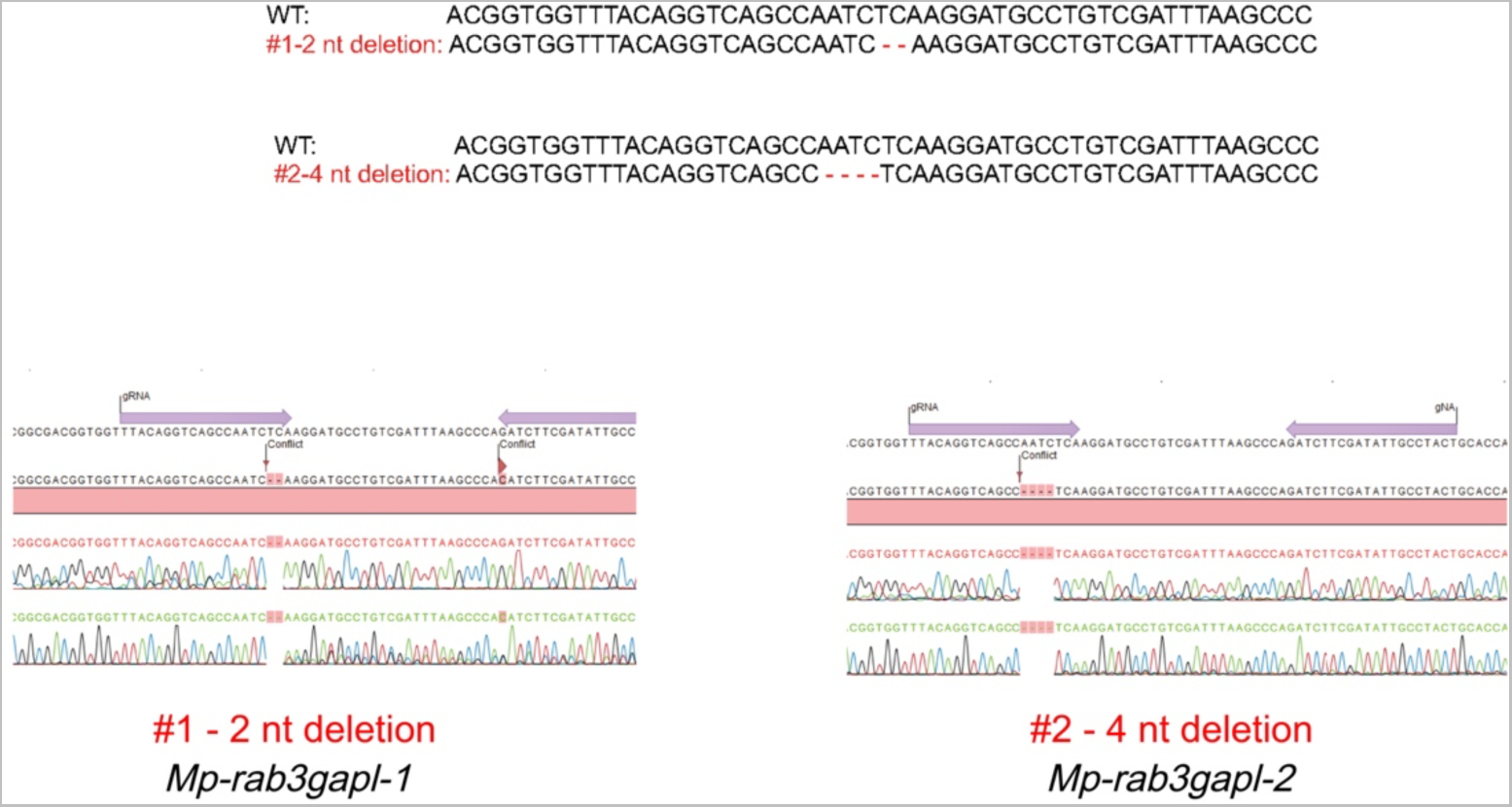
Generation of two independent *M. polymorpha Rab3GAPL* CRISPR knockout mutants, designated as *Mp-rab3gapl-1* and *Mp-rab3gapl-2*, in a GFP:ATG8b background. The top panel illustrates a comparison between wild-type (WT) and *Mp-rab3gapl-1* and *-2* mutations. The bottom panel displays chromatograms depicting the sequencing results of the WT and mutant plants.

**Figure S3F.**
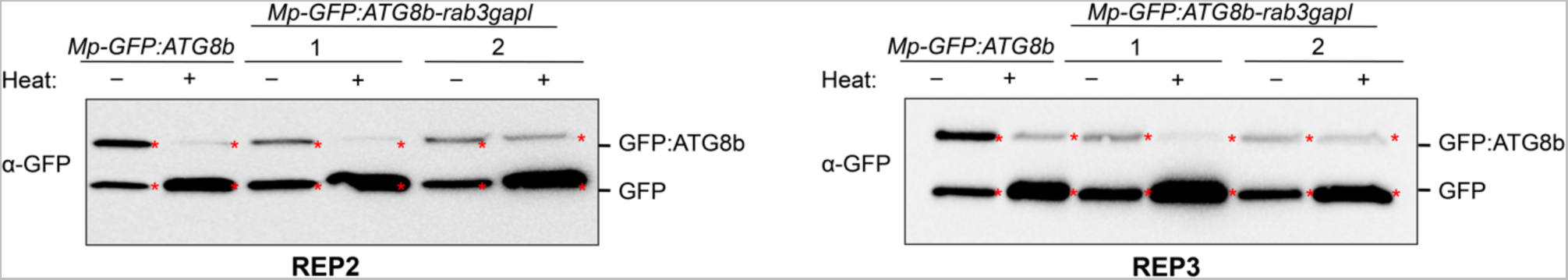
*Marchantia polymorpha* Rab3GAPL-KO mutants have increased ATG8 autophagic flux (additional repeats for Figure 3D). Autophagic flux analysis of WT and Rab3GAPL-KO mutants in MpEF::GFP:ATG8b background after 6 hours of heat stress treatment following 2 hours recovery. Flux is estimated as a measure of ratio between free GFP to full size GFP:ATG8b. Both Rab3GAPL-KO mutants showed increased GFP/GFP:ATG8b protein signal ratio under heat stress and control conditions compared to the control plants. Protein extracts were prepared using 14-day-old thalli and immunoblotted.

**Figure S4A.**
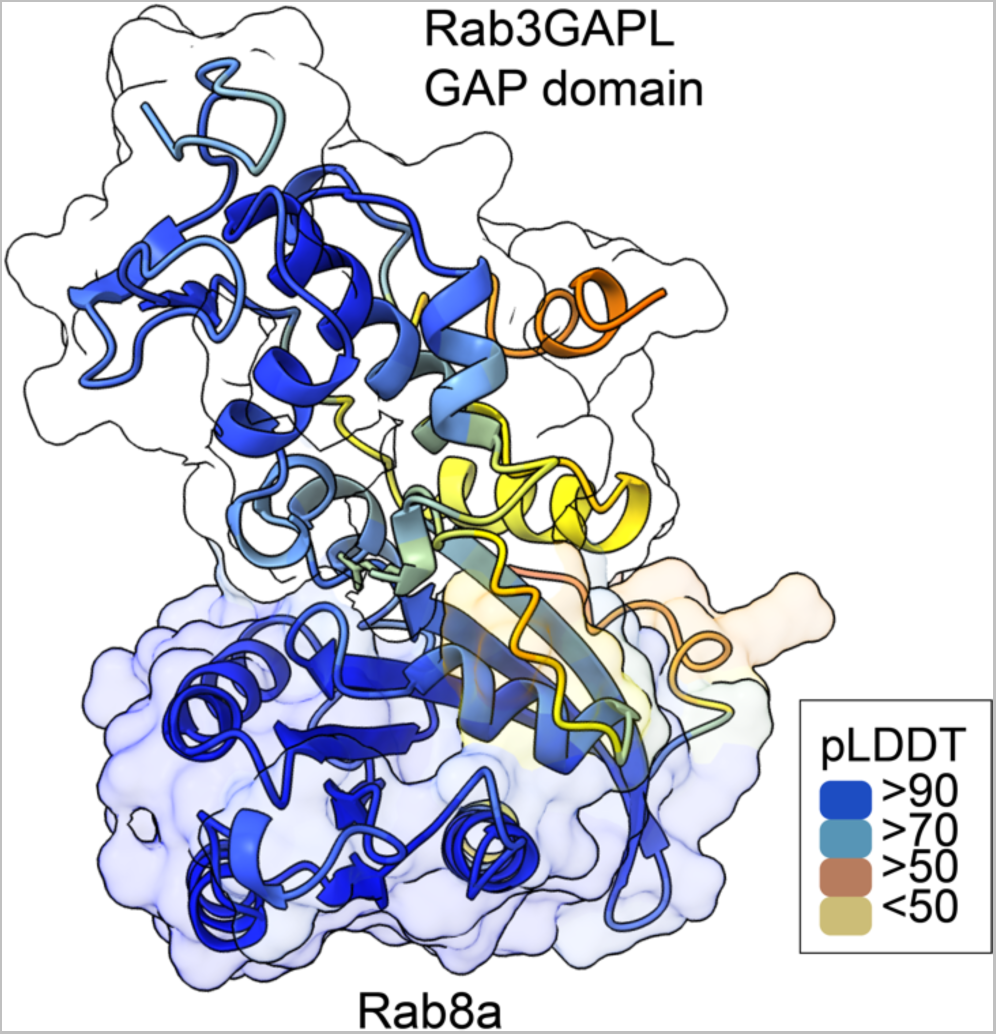
AF2-M predicted model of Rab8a in complex with Rab3GAPL GAP domain. The colors of Rab3GAPL GAP domain and Rab8a represent the confidence score (pLDDT) calculated by AF2, as indicated in the rectangular box.

**Figure S4B.**
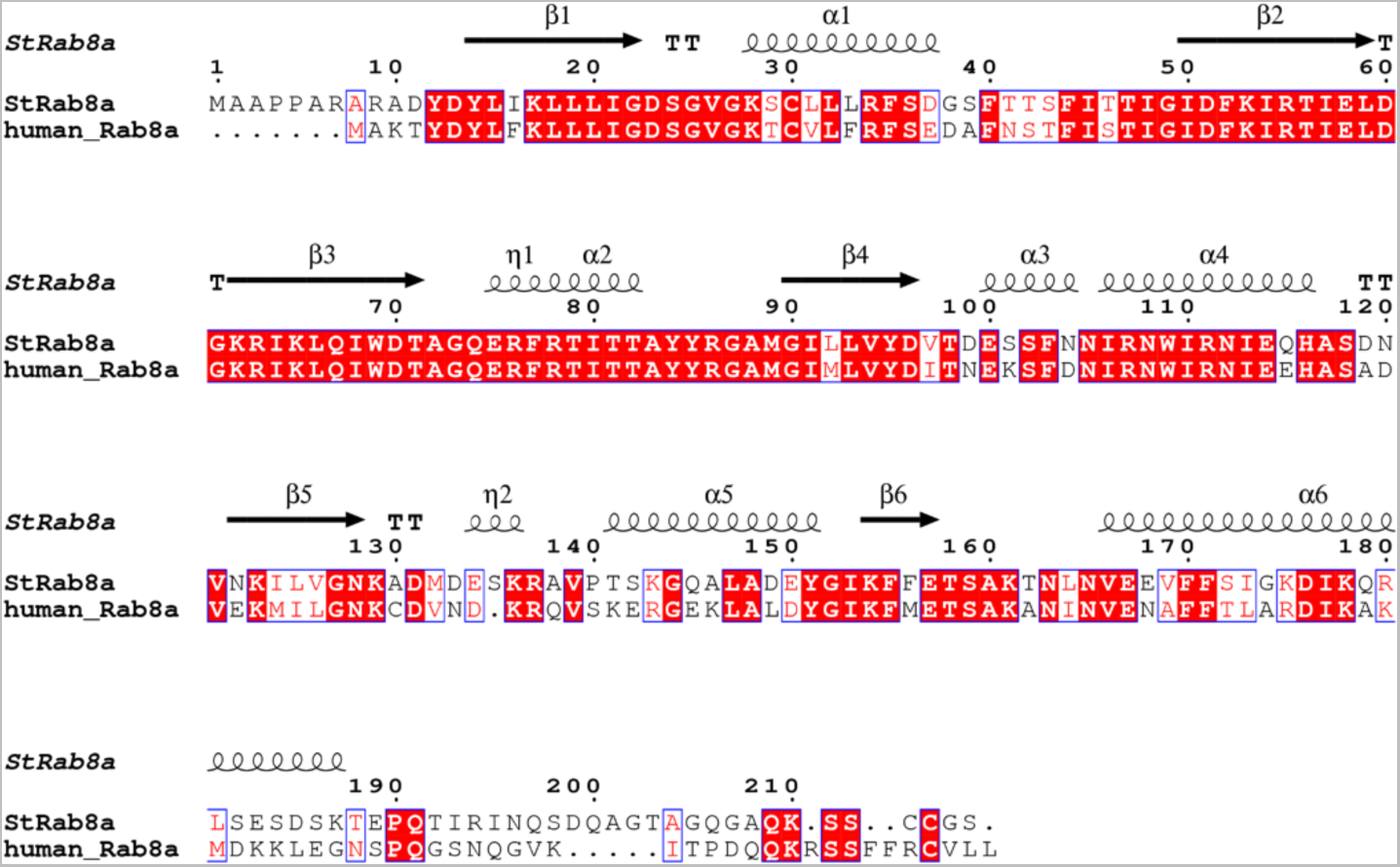
Pairwise sequence alignment of potato Rab8a and human Rab8a. Alignments were obtained using the MUSCLE algorithm and were visualized and color-coded via ESPript 3.0 (*39*). The Rab8a displayed a high degree of protein sequence conservation with 68% amino acid identity.

**Figure S4C, D.**
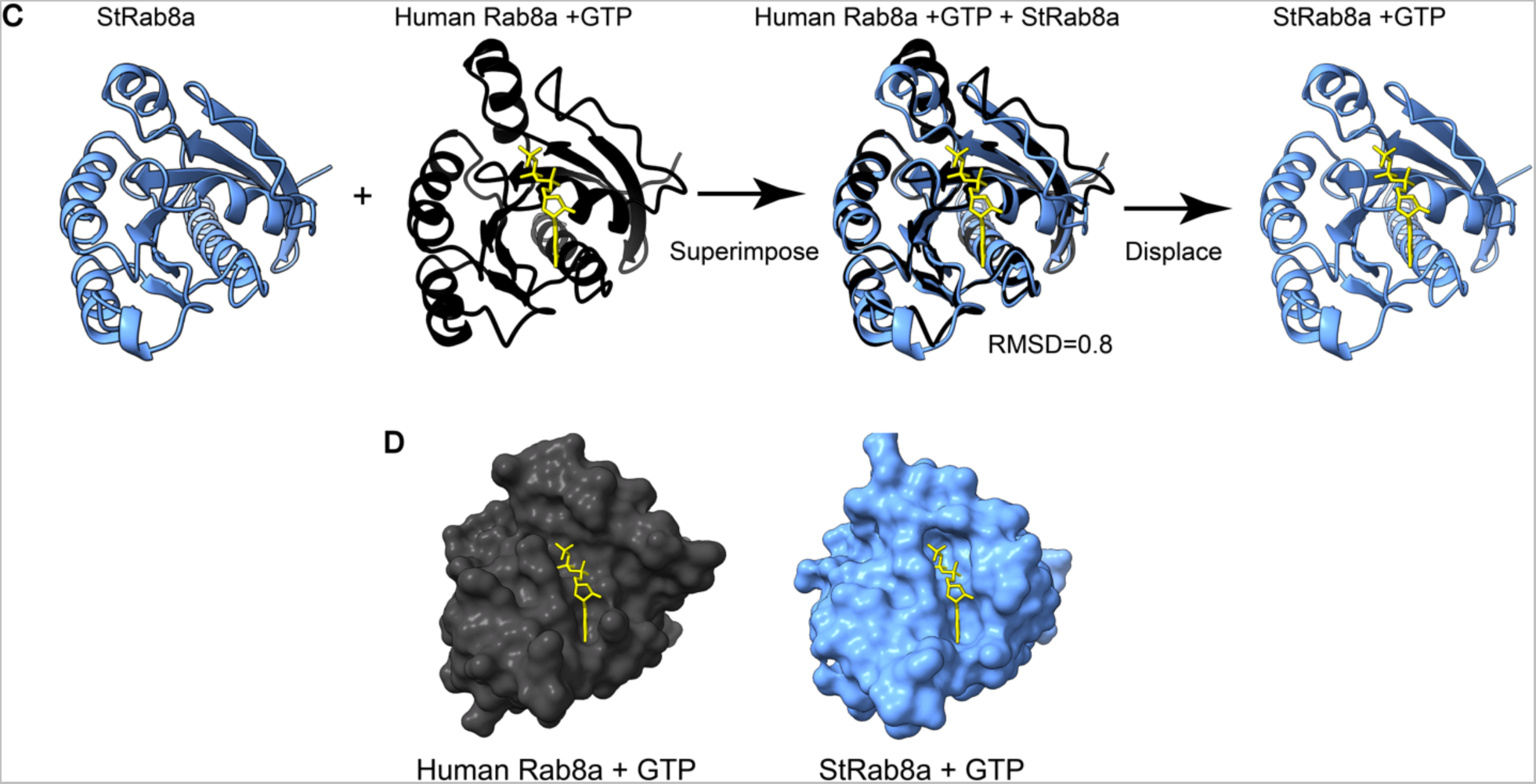
AF2-guided *ab initio* molecular replacement of GTP-bound human Rab8a with the potato Rab8a. **(C)** Superposition of experimentally determined structure of human Rab8a (black) bound to GTP (yellow) (PDB:6WHE) with AF2 prediction of potato Rab8a (blue) structure. Following superimposition, human Rab8a structure was removed, resulting in potato Rab8a bound to GTP in the nucleotide binding pocket. **(D)** Side-by-side surface views of human Rab8a (PDB:6WHE) and potato Rab8a AF2-model with GTP-bound conformations.

**Figure S4E.**
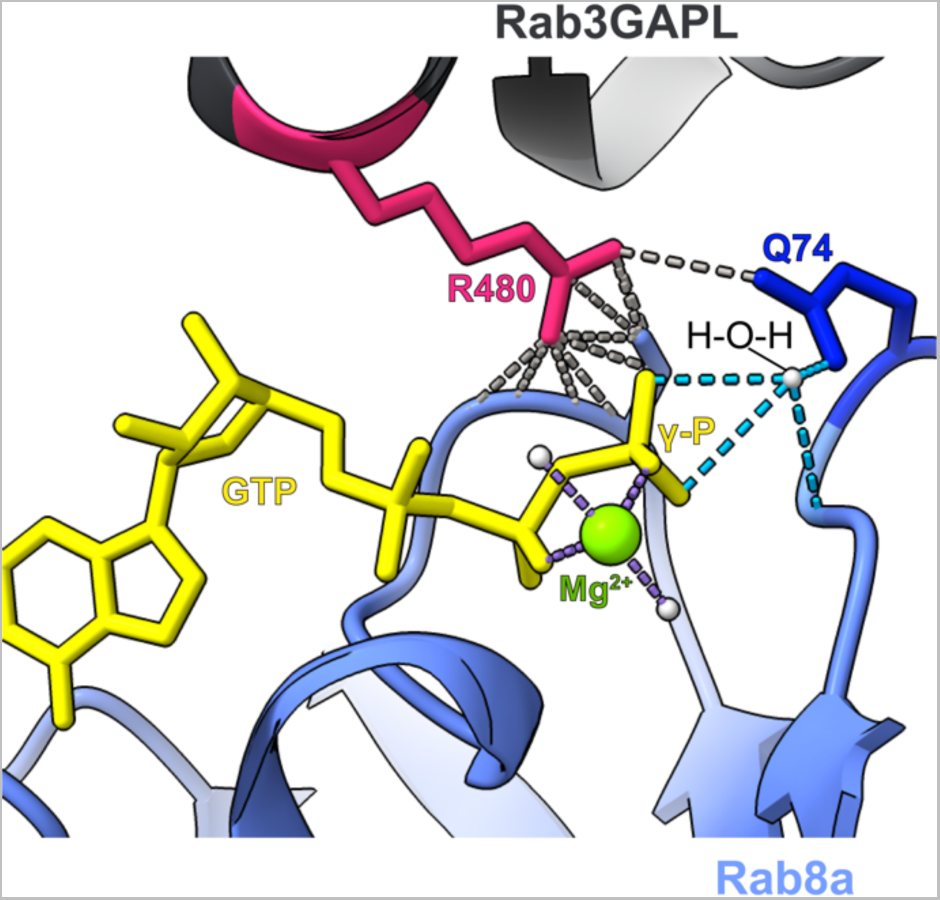
Predicted AF2-M model of Rab8a in complex with the GAP domain of Rab3GAPL. The catalytic arginine residue, R480 (magenta), of Rab3GAPL is located in close proximity to the GTP binding pocket of Rab8a. R480 forms contacts with the switch-2 glutamate (Q74, depicted in dark blue) of Rab8a. Q74 forms hydrogen bonds with the nucleophilic water molecule that act on gamma-phosphate (γ-P) of the GTP molecule (yellow).

**Figure S5A, B.**
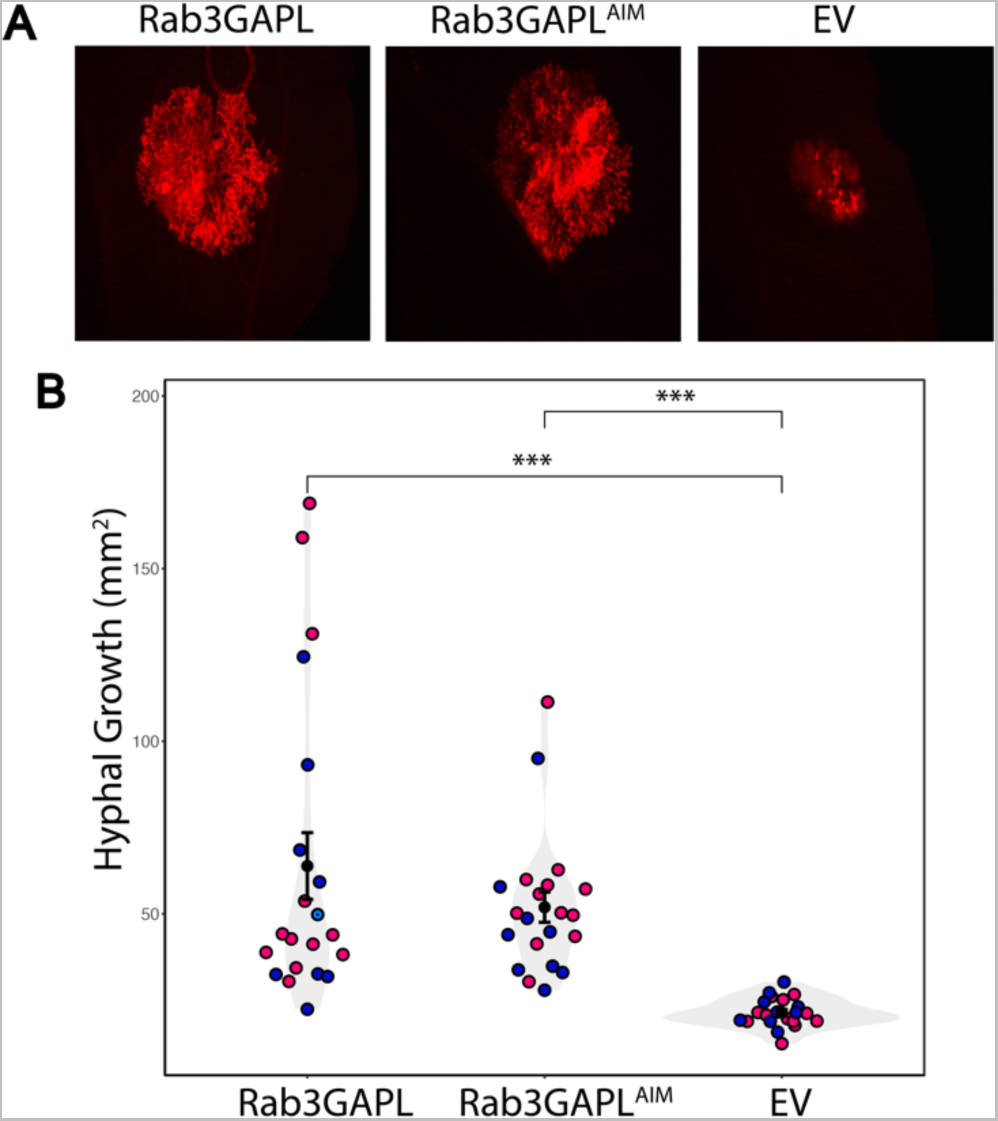
Rab3GAPL increases hyphal growth of *P. infestans* in an AIM-independent manner. (A) *N. benthamiana* leaves expressing Rab3GAPL, Rab3GAPL^AIM^ or EV control were infected with tdTomato-expressing *P. infestans*, and pathogen growth was calculated by measuring hyphal growth of pathogen using fluorescence stereomicroscope at 5 days post infection. (B) Both Rab3GAPL expression (63.9, N = 21 spots) and Rab3GAPL^AIM^ expression (51.9, N = 21 spots) significantly increases *P. infestans* hyphal growth compared to EV control (21.4, N = 21 spots). Statistical differences were analyzed by Mann-Whitney U test in R. Measurements were highly significant when p<0.001 (***).

**Figure S5C, D.**
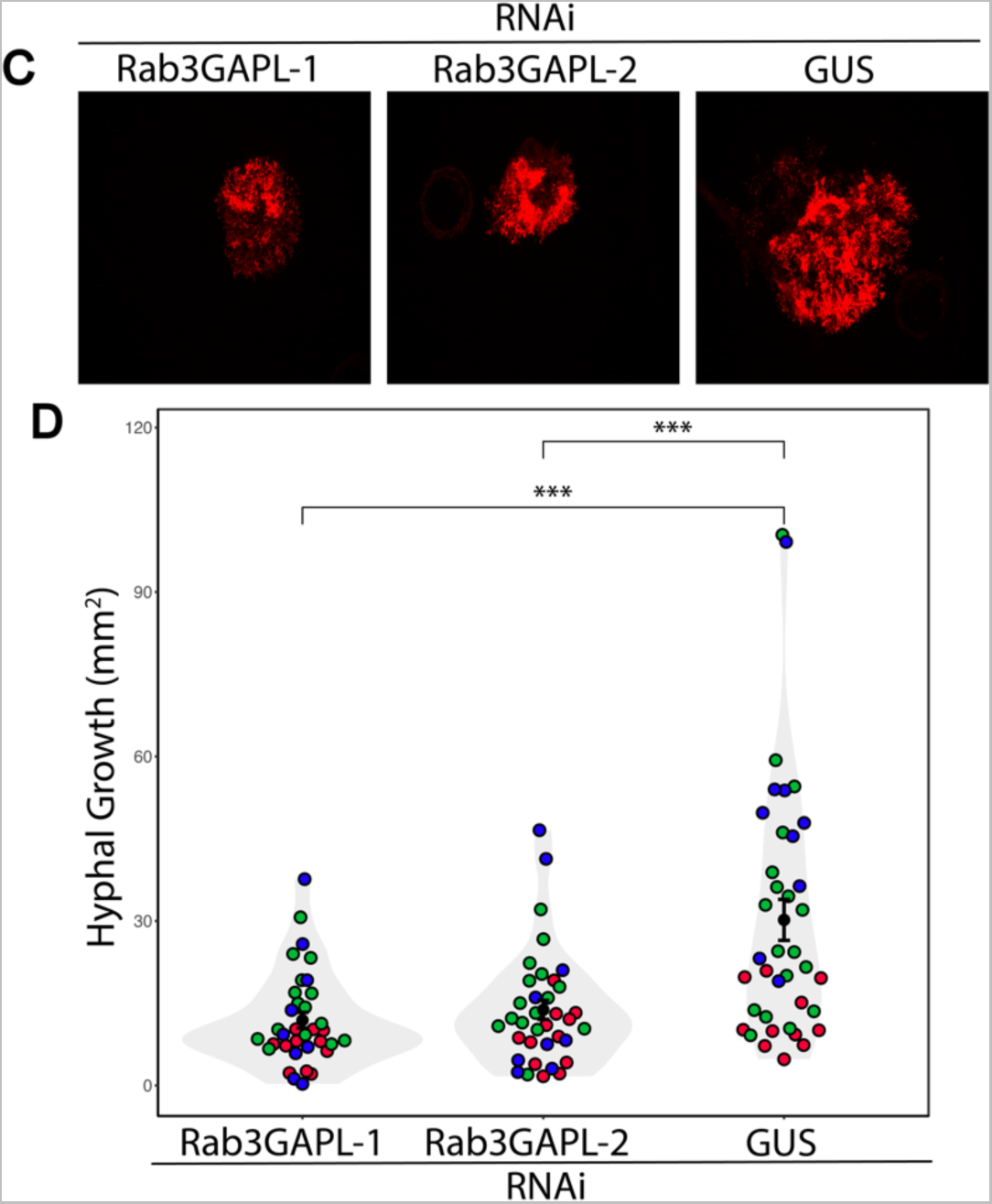
Silencing *Rab3GAPL* significantly reduces hyphal growth of *P. infestans*. (C) *N. benthamiana* leaves expressing RNAi:Rab3GAPL-1, RNAi:Rab3GAPL-2 or RNAi:GUS control were infected with tdTomato-expressing *P. infestans*, and pathogen growth was calculated by measuring hyphal growth of pathogen using fluorescence stereomicroscope at 5 days post infection. (D) Both RNAi:Rab3GAPL-1 expression (11.9, N = 35 spots) and RNAi:Rab3GAPL-2 expression (13.8, N = 36 spots) significantly reduces *P. infestans* hyphal growth compared to RNAi:GUS control (30.2, N = 38 spots). Statistical differences were analyzed by Mann-Whitney U test in R. Measurements were highly significant when p<0.001 (***).

**Table S1. Primers used in this work**

**Table S2. Details of constructs used**

**Table S3. Proteins and sequences used for AF2**

**Table S4. Details of antibodies used**

**Table S5. Summary of statistics**

## Acknowledgments

We acknowledge and thank the Imperial College FILM facility for their technical expertise and the use of their microscopy equipment. We would also like to thank Vienna Biocenter Core Facilities for plant sciences, bio-optics, molecular biology, and protein technology core facilities. We also express our gratitude to the members of the Bozkurt, Dagdas and Kamoun groups for their suggestions and productive discussion.

## Author contributions

Conceptualization: T.O.B., Y.D. Methodology: E.L.H.Y., A.Y.L., M.C., A.M., L.P., T.O.B. Validation: E.L.H.Y., A.Y.L., M.C., A.M., L.P. Formal Analysis: E.L.H.Y., A.Y.L., M.C., Y.T., A.M., L.P. Investigation: E.L.H.Y., A.Y.L., M.C., Y.T., A.M., L.P., M.J., P.P., C.D., T.O.B. Data Curation: E.L.H.Y., T.O.B. Visualization: E.L.H.Y., A.Y.L., M.C., A.M., L.P., T.O.B. Writing – Original Draft: E.L.H.Y., A.Y.L., T.O.B. Writing – Review & Editing: E.L.H.Y., Y.D.,T.O.B. Supervision: C.D., E.C., Y.D., T.O.B. Funding Acquisition: Y.D., T.O.B.

## Data and Materials Availability

All relevant study data are included in the article, and in the Supplementary Materials files. AF2- multimer predictions are uploaded to the public repository Figshare and is available at https://doi.org/10.6084/m9.figshare.23587575.

